# Mrx6 binds the Lon protease Pim1 N-terminal domain to confer selective substrate specificity and regulate mtDNA copy number

**DOI:** 10.1101/2025.02.14.638270

**Authors:** Simon Schrott, Charlotte Gerle, Sebastian Bibinger, Lea Bertgen, Giada Marino, Serena Schwenkert, Romina Rathberger, Johannes M. Herrmann, Christof Osman

**Affiliations:** Faculty of Biology, Ludwig-Maximilians-Universität München, Großhaderner Straße 2 - 4, Planegg-Martinsried 82152, Germany; Cell Biology, University of Kaiserslautern, RPTU, Erwin-Schrödinger-Strasse 13, 67663 Kaiserslautern, Germany

## Abstract

Mitochondrial DNA (mtDNA) copy number regulation remains incompletely understood, despite its importance in cellular function. In *Saccharomyces cerevisiae*, Mrx6 belongs to the Pet20-domain-containing protein family, consisting of Mrx6, Pet20, and Sue1. Notably, absence of Mrx6 leads to increased mtDNA copy number. Here, we identify the C-terminus of Mrx6 as essential for its stability and interaction with the mitochondrial matrix protein Mam33. Deletion of Mam33 mimics the effect of Mrx6 loss, resulting in elevated mtDNA copy number. Bioinformatics, mutational analyses, and immunoprecipitation studies reveal that a subcomplex of Mam33 and Mrx6 trimers interacts with the substrate recognition domain of the conserved mitochondrial Lon protease Pim1 through a bipartite motif in the Pet20 domain of Mrx6. Loss of Mrx6, its paralog Pet20, Mam33, or mutations disrupting the interaction between Mrx6 and Pim1 stabilize key proteins required for mtDNA maintenance, the RNA polymerase Rpo41 and the HMG-box-containing protein Cim1. We propose that Mrx6, alongside Pet20 and Mam33, regulates mtDNA copy number by modulating substrate degradation through Pim1. Additionally, Mrx6 loss alters Cim1’s function, preventing the detrimental effect on mtDNA maintenance observed upon Cim1 overexpression. The presence of three Pet20-domain proteins in yeast implies broader roles of Lon protease substrate recognition beyond mtDNA regulation.

## INTRODUCTION

Mitochondria contain their own genome (mtDNA) that encodes crucial subunits required for oxidative phosphorylation. mtDNA maintenance is therefore of critical importance for cellular health and energy supply. Multiple copies of mtDNA are present within the mitochondrial network and alterations in mtDNA copy number (mtDNA CN) have been linked to numerous diseases and aging (Gupta et al., 2023; Filograna et al., 2021; Castellani et al., 2020). However, the cellular mechanisms that regulate mtDNA CN are incompletely understood.

In *Saccharomyces cerevisiae*, mtDNA CN scales with mitochondrial network length and cell size (Osman et al., 2015; Seel et al., 2023). We recently conducted a large-scale analysis to identify genes that, when absent, lead to altered mtDNA CN (Göke et al., 2020). Among the candidates were the genes *CIM1* and *MRX6*, the loss of which results in increased mtDNA CN. *CIM1* encodes a mtDNA-binding HMG-box containing protein with similarities to the well-characterized mtDNA packaging proteins Abf2 in yeast and TFAM in mammals (Schrott & Osman, 2023). While absence of Cim1 increases mtDNA CN, its overexpression leads to defects in mtDNA maintenance, which has led to the hypothesis that Cim1 negatively regulates mtDNA replication.

Mrx6 belongs to a protein family, whose members are defined by a poorly characterized Pet20 domain (Göke et al., 2020). In *S. cerevisiae*, the Pet20 domain protein family comprises three members: Mrx6, Pet20, and Sue1. While the role of Mrx6 in regulating mtDNA copy number remains unclear, it has been identified as a key interaction partner of the conserved Lon protease Pim1 (Göke et al., 2020; Michaelis et al., 2023; Bertgen et al., 2024). This interaction suggests that Mrx6 may facilitate the degradation of proteins involved in mtDNA replication (Göke et al., 2020). Furthermore, Mrx6 associates with Pet20 and a protein called Mam33 (Göke et al., 2020; Hillman & Henry, 2019). If and how these proteins function together to regulate mtDNA CN has not been analyzed.

Lon proteases are hexameric assemblies that are conserved from bacteria to humans and different domains can be distinguished (Yang et al., 2022). The C-terminal parts contain an ATPase and a protease domain, which are required for unfolding and degradation of substrates, respectively. The function of the N-terminal domain is less well studied, but has been implicated in substrate selection (Li et al., 2021). The Lon protease has been linked to mtDNA CN control in mammals, flies and *S. cerevisiae*. Studies suggested that the Lon protease degrades the mtDNA packaging factor TFAM and thereby influences mtDNA packaging and in turn mtDNA CN (Lu et al., 2013; Matsushima et al., 2010). In *S. cerevisiae* it was recently shown that Pim1 is involved in degradation of the Abf2 paralog Cim1, whereas Abf2 does not appear to be a strong substrate of Pim1 *in vivo* (Schrott & Osman, 2023). Since Cim1 overexpression results in mtDNA maintenance defects, efficient degradation of Cim1 by Pim1 was proposed to be important for mtDNA replication. Recently, a comprehensive genome-wide association study has linked mutations in the human Pim1 homolog LonP to altered mtDNA levels in blood samples (Gupta et al., 2023). Hence, the Lon protease emerges as a central player in mtDNA CN regulation from yeast to humans. An outstanding question concerns the substrate specificity of the Lon protease. On one hand it has been proposed that Lon degrades misfolded and damaged proteins through recognition of unfolded protein stretches (Janowsky et al., 2005; Wagner et al., 1994). On the other hand specific degradation of factors for regulatory purposes has been proposed (Göke et al., 2020; Schrott & Osman, 2023; Langklotz & Narberhaus, 2011; Jonas et al., 2013).

Here, we examine the interplay between Mrx6, Mam33, Pet20 and Pim1 to understand their role in mtDNA CN regulation. We find that Mrx6, Mam33 and likely Pet20 form a subcomplex that sits like a gatekeeper on the N-termini of the hexameric Pim1 complex. Absence or depletion of any of these subunits results in an accumulation of the RNA polymerase Rpo41 and the protein Cim1. These results suggest that Mrx6, Mam33 and Pet20 regulate degradation of proteins by the Lon protease and thereby regulate mtDNA CN.

## RESULTS

### Mrx6-Dependent Interactions of Pim1 with Mam33 and Pet20

We have previously found that Mrx6 interacts with Pim1, Mam33 and Pet20 through immunoprecipitation of Mrx6- or Pet20-3xFlag (Göke et al., 2020). To better understand the architecture of potential complexes, we performed immunoprecipitation experiments of Spot-tagged® Pim1 from mitochondria of cells containing or lacking Mrx6. The Spot-tag, composed of 12 amino acids, was optimized from the BC2-tag system for enhanced high-affinity nanobody binding (Traenkle et al., 2015; Braun et al., 2016; Virant et al., 2018). Expression of endogenously Spot-tagged Pim1 did not affect mtDNA CN (Figure 1A). In cells expressing Pim1-Spot, the deletion of *MRX6* resulted in increased mtDNA CN, typical for cells lacking Mrx6 (Göke et al., 2020). Moreover, Pim1-Spot strains did not exhibit elevated frequencies of petite formation or growth defects on both fermentable and non-fermentable media (Figure 1B and Supplementary Figure S2A). These findings indicate that Spot-tagged Pim1 is functional.

**Figure 1.**
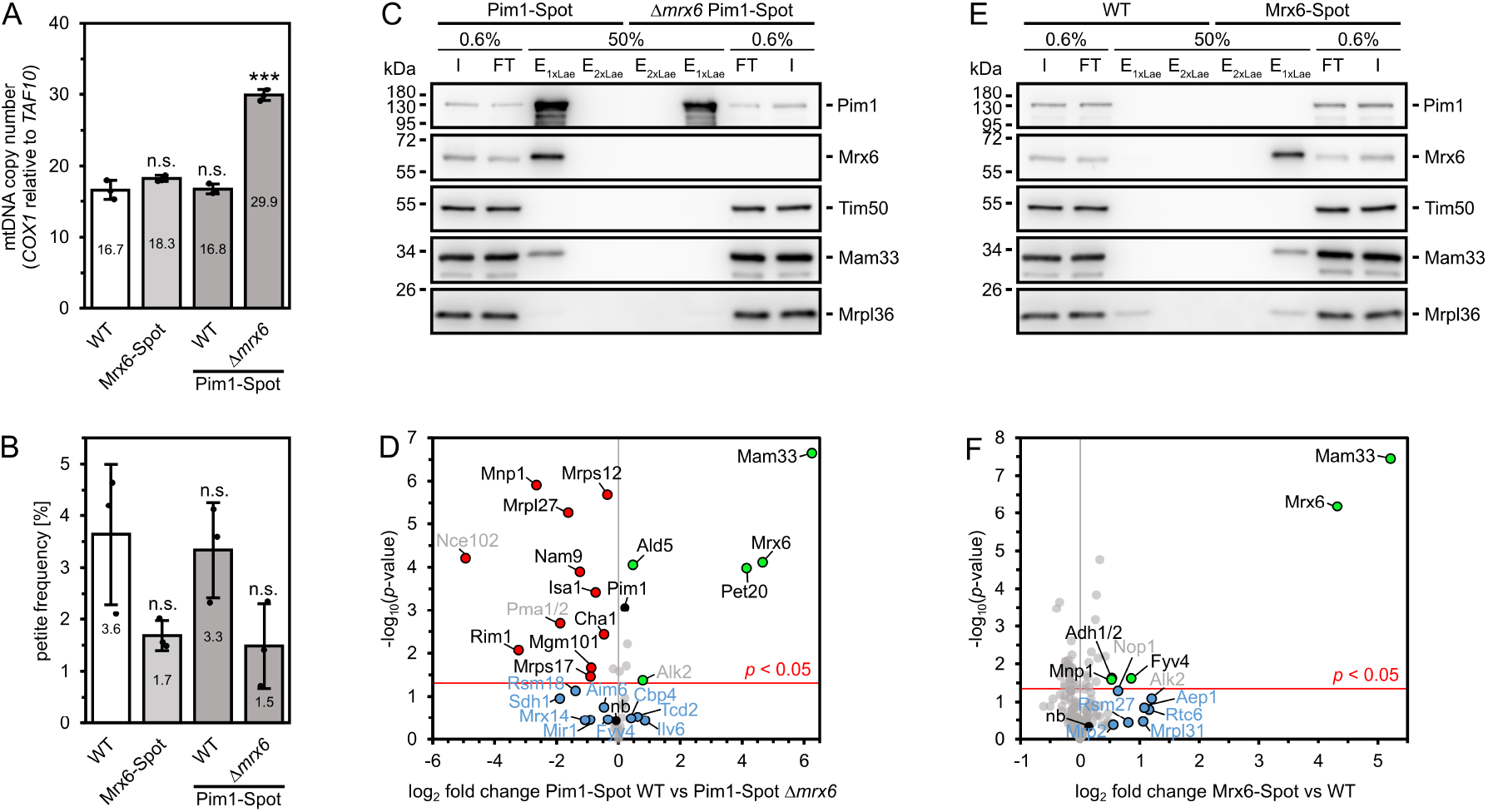
Mrx6-dependent interactions of Pim1 with Mam33 and Pet20. (**A**) qPCR on mtDNA-encoded *COX1* and nuclear reference *TAF10* of indicated strains grown in YPD at 30°C. n = 3, data represent mean ± SD. (**B**) Petite frequency of indicated strains grown in YPD at 30°C. n = 3, data represent mean ± SD. (**C+E**) Immunoblots of Pim1, Mrx6, Tim50 (control), Mam33, and Mrpl36 from Spot-tag immunoprecipitations using magnetic particle beads and lysed mitochondria isolated from Pim1-Spot and Δ*mrx6* Pim1-Spot (C) or WT and Mrx6-Spot (E) cells, cultivated in YPG at 30°C. Input (I) and flow-through (FT) fractions represent 0.6% of total amounts and bound fractions eluted with 1x (E_1xLae_) followed by 2x Laemmli buffer (E_2xLae_) are 50% of eluted volumes. (**D+F**) Volcano plots of liquid chromatography-tandem mass spectrometry (LC-MS/MS) detected proteins after on-bead digest of Spot-tag immunoprecipitations using magnetic particle beads and lysed mitochondria isolated from Pim1-Spot and Δ*mrx6* Pim1-Spot (D) or WT and Mrx6-Spot (F) cells cultivated in YPG at 30°C. n = 3 (technical), data was normalized by LFQ. Significantly enriched/decreased hits are highlighted in green/red. Labels for non-mitochondrial proteins are colored in gray. Putative interesting but not significant hits with a log_2_ fold change > 0.3 (D) or > 0.4 (F) are colored in blue. “nb” shows BC2 nanobody abundance. Pim1-Spot is highlighted in D.

In line with a prior high-throughput analysis involving Pim1-GFP (Michaelis et al., 2023), immunoprecipitation of Pim1-Spot from mitochondrial lysates with anti-Spot-tag nanobody followed by immunoblots revealed the co-purification of Mrx6 and Mam33, irrespective of bead type or protease inhibitor used (Figure 1C and Supplementary Figure S2B). Strikingly, no interaction between Pim1 and Mam33 was observed when Pim1-Spot was purified from Δ*mrx6* mitochondria. To gain a comprehensive understanding of proteins that interact with Pim1 in a Mrx6-dependent manner, we performed liquid chromatography-tandem mass spectrometry (LC-MS/MS) analysis on proteins purified with Pim1-Spot from either wild-type (WT) or Δ*mrx6* mitochondria. These analyses confirmed that the co-purification of Mam33 with Pim1-Spot depends on the presence of Mrx6 (Figure 1D). Additionally, these analyses demonstrated that the Mrx6-homolog Pet20 was identified only in Pim1-Spot purifications from WT mitochondria, but not from Δ*mrx6* mitochondria, indicating that a stable interaction between Pet20 and Pim1 relies on Mrx6. Interestingly, the third Pet20 domain containing protein, Sue1, interacted with Pim1 independently of Mrx6 and even co-purified with Pim1 more efficiently in the absence of Mrx6 (LFQ ~2- and iBAQ ~3-fold more, Supplementary Tables 2).

While Pet20 and Mam33 co-purify with Pim1 exclusively in the presence of Mrx6, distinct proteins are enriched with Pim1-Spot specifically in Δ*mrx6* cells. These proteins include mitochondrial ribosomal components from both the small (Nam9, Mrps17, and Mrps12) and large subunits (Mnp1 and Mrpl27), as well as mtDNA-associated proteins such as Mnp1, Rim1, Mgm101, and Cha1. The underlying cause for the preferential interaction of these proteins with Pim1 in the absence of Mrx6 remains to be elucidated. Nonetheless, it is plausible that Mrx6 modulates the activity or binding affinity of Pim1 to its substrates.

To further investigate the interaction network between Pim1, Pet20 domain proteins, and Mam33 from the perspective of Mrx6, we performed immunoprecipitations using Spot-tagged Mrx6. Mrx6-Spot strains show mtDNA levels akin to WT but exhibit a reduced petite frequency, similar to Δ*mrx6* strains, suggesting compromised Mrx6 functionality (Figures 1A+B). Notably, Mrx6-Spot immunoprecipitation and subsequent analysis of co-purified proteins via immunoblots and LC-MS/MS revealed a selective enrichment of Mam33 but did not co-purify Pim1 or Pet20 (Figures 1E+F and Supplementary Figure S2C). This contrasts with our prior observation in which, alongside Mam33, both Pim1 and Pet20 co-purified with Mrx6-3xFlag (Göke et al., 2020). We ascribe this discrepancy to the Spot-tag, which may impede stability of the Pim1-Mrx6 interaction compared to the Flag-tag. The decreased petite frequency of a strain expressing Mrx6-Spot aligns with a compromised function of Mrx6-Spot. Crucially, the co-purification experiment indicated that the interaction between Mrx6-Spot and Mam33 is independent of Pim1.

Taken together, our results reveal that Mrx6 is essential for a stable interaction between Pim1 and Pet20 as well as Mam33. Our finding that the functionally compromised Mrx6-Spot interacts with Mam33, but not Pim1 or Pet20, further suggests that Mrx6 and Mam33 constitute a stable subcomplex linked to Pim1 through Mrx6.

### Interdependency of Mrx6 and Mam33 governs mitochondrial DNA copy number

We next focused on understanding how Mrx6 interacts with Mam33. The Mrx6 protein is comprised of two distinct segments. The N-terminal region of Mrx6 encompasses a Pet20 domain, which is similarly present in its homologs Pet20 and Sue1. In contrast, Mrx6 also possesses a unique C-terminal segment that is absent in Pet20 and Sue1 (Figure 2A and Supplementary Figures S1A-C). AlphaFold predictions indicated a structurally robust folding for the C-terminal domain of Mrx6, in contrast to its N-terminal Pet20 domain, which showed lower structural organization and confidence scores (Figures 2B-D). To explore the potential interaction interfaces between Mrx6 and Mam33, we performed Alphafold predictions with three Mam33 molecules due to its known trimeric configuration (Pu et al., 2011), excluding its mitochondrial targeting sequence (MTS) (1-46) (Vögtle et al., 2009), and a single Mrx6 molecule devoid of its MTS (1-33). This analysis revealed a high-confidence interaction of the positively charged C-terminus of Mrx6 and the negatively charged, bowl-shaped cavity of the Mam33 trimer (Supplementary Figures S3A-E). Conversely, the Pet20 domain of Mrx6 did not appear to be involved in this interaction.

**Figure 2.**
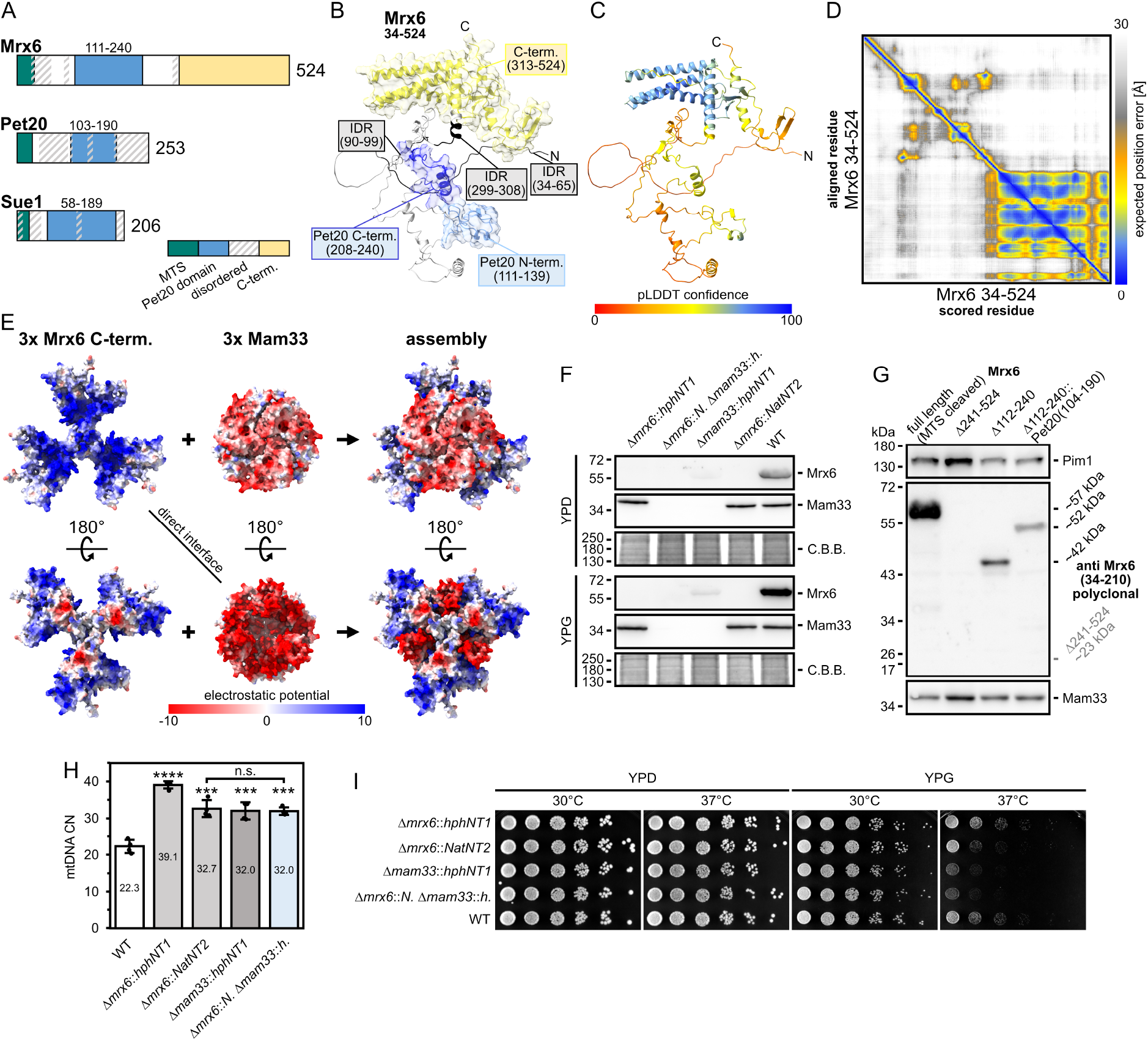
Interdependency of Mrx6 and Mam33 governs mitochondrial DNA copy number. (**A**) Schematic of protein domains of Mrx6, Pet20 and Sue1. Disordered regions (flDPnn propensity (Hu et al., 2021)) are highlighted diagonal lines. (**B-D**) Highest ranked (by predicted Template Modeling score - pTM) AlphaFold monomer prediction of Mrx6 (residues 34-524) lacking the mitochondrial targeting sequence (MTS) (mean pLDDT confidence ~59.74). The ribbon model shows the N_Pet20_- and N_Pet20_-motifs in blue and the C-terminal region in yellow. Intrinsically disordered regions (IDR) are colored in black (B). Error estimates are either shown as ribbon structure colored according to the residue specific pLDDT confidence (C) or as a heat map plot of the expected position error for every pair of residues (D). (**E**) AlphaFold multimer prediction of 3x Mam33 (residues 47-266) and 3x Mrx6 C-terminus (residues 271-524). Best ranked model with improved pTM + pTM confidence of ~0.7674. Surfaces are colored according to the coulombic electrostatic potential in the range of −10 to 10. (**F+G**) Immunoblots of indicated strains grown in YPD or YPG (G only YPD) at 30°C with Mrx6 and Mam33 (Pim1 additional loading control in G, for more controls of F see 4D) antibodies. C.B.B. stained protein bands remaining in the gel after transfer served as a loading control. n = 1. (**H**) qPCR on mtDNA encoded *COX1* and nuclear reference *TAF10* of indicated strains grown in YPD at 30°C. n = 4, each mutant 2 biological replicates from two independent deletion strains, data represent mean ± SD. (**I**) Drop dilution growth analysis at 30 or 37°C of indicated strains pre-grown in YPD at 30°C. Images were taken after 48 h growth on YPD or YPG.

LC-MS/MS analysis of the Pim1 complex revealed Mrx6 presence at an average ratio of ~1:2.2 relative to Mam33 (even ~1:1.3 in data by Bertgen et al.), rather than a 1:3 ratio (Supplementary Tables 2). Thus, we postulated that more than one molecule of Mrx6 might interact with the trimeric Mam33. This idea is further supported by the structural symmetry observed in the predictions, where Mrx6 interacts with the Mam33 trimer at the interface of two adjacent Mam33 subunits. Indeed, a high-confidence prediction indicated that the trimer of Mam33 can accommodate up to three C-terminal regions of Mrx6, leading to a compact structural formation (Figure 2E and Supplementary Figures S3F-J). In agreement with the distinctive C-terminal domain of Mrx6, no interactions were predicted between Mam33 and either Pet20 or Sue1. Furthermore, no high-confidence interactions were observed between any domain of Pim1 and Mam33.

The resulting compact structure, composed of three Mrx6 and three Mam33 molecules, led us to ask if absence of either protein affects the stability of the other. While Mam33 levels remained unaltered in Δ*mrx6* cells, Mrx6 was virtually undetectable in Δ*mam33* cells (Figure 2F). Similarly, a construct of Mrx6 lacking the C-terminal domain, predicted to bind Mam33, did not accumulate to detectable levels (Figure 2G). In contrast, Mrx6 variants either lacking the Pet20-domain or incorporating the Pet20 domain from the Pet20 protein accumulated to detectable but lower levels. These results suggest that the stability of Mrx6 depends on its binding to Mam33 and that both proteins form a functional unit.

Since lack of Mrx6 results in increased mtDNA CN, we assessed whether deletion of Mam33 would similarly increase mtDNA CN. Indeed, mtDNA levels were elevated ~1.5-fold in Δ*mam33* cells, comparable to the increase observed in Δ*mrx6* cells (Figure 2H and Supplementary Figures S3N-P). No further rise in mtDNA levels was observed in Δ*mam33*Δ*mrx6* double mutants. Notably, an analogous increase in mtDNA CN was observed in cells expressing a Mrx6 truncation, lacking its C-terminal extension (Figure 3G). We further compared growth of Δ*mam33*, Δ*mrx6* and Δ*mam33*Δ*mrx6* mutant strains, which was virtually indistinguishable from WT on fermentable medium (Figure 2I and Supplementary Figures S3K-M). On medium containing a non-fermentable carbon source, we observed growth defects of Δ*mam33* and Δ*mrx6* single mutants at 37°C, which was more pronounced in Δ*mam33* cells. The growth of the Δ*mam33*Δ*mrx6* double mutant closely mirrored that of Δ*mam33* cells, thus revealing that *MAM33* is epistatic over *MRX6*. Lastly, the low petite frequency observed in Δ*mrx6* cells was also phenocopied in Δ*mam33* and Δ*mrx6*Δ*mam33* cells, further supporting the idea that both proteins function together (Supplementary Figure S3Q).

**Figure 3.**
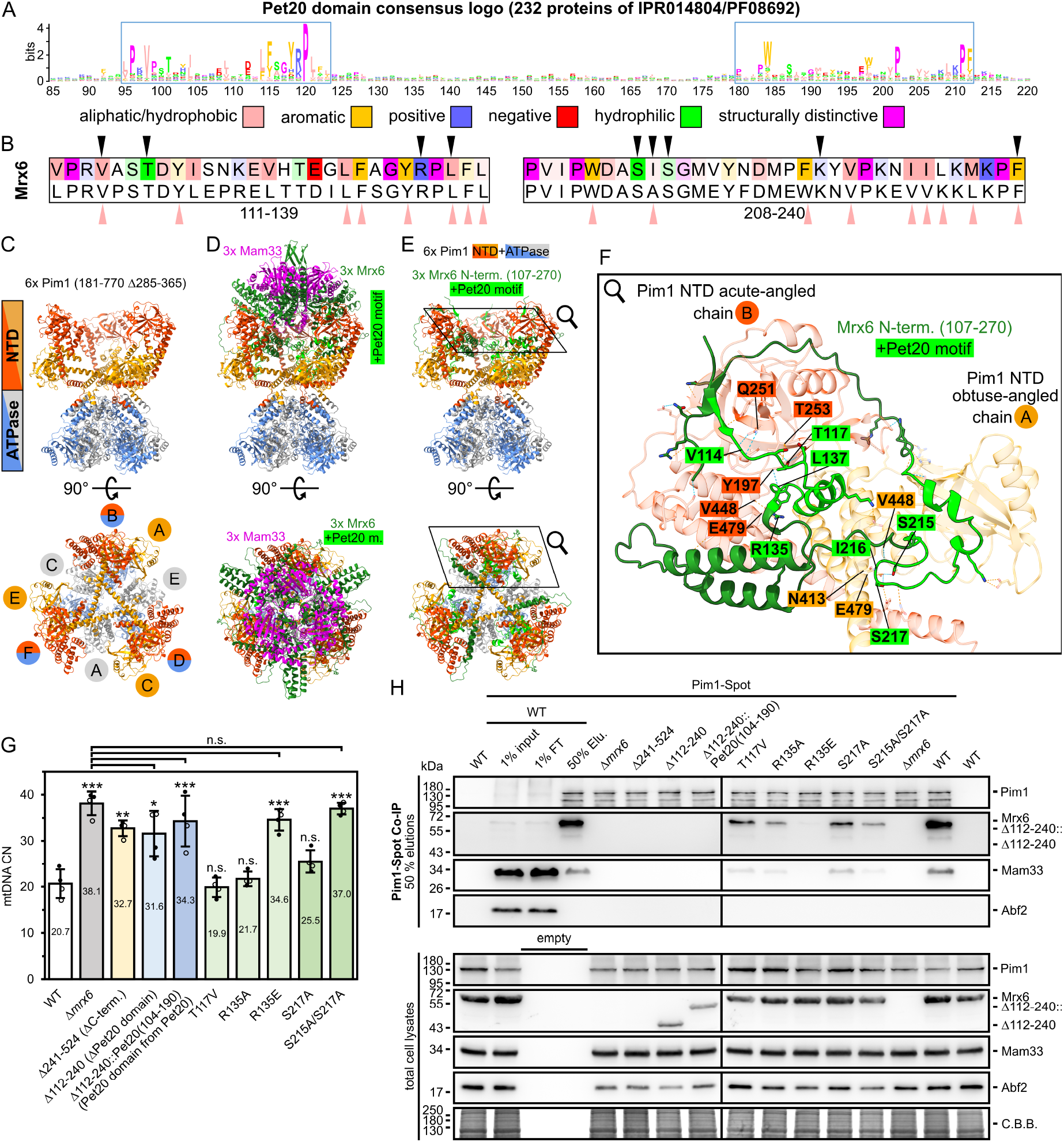
A bipartite-motif in the Pet20 domain mediates interaction between Mrx6 and Pim1. (**A**) Pet20 domain consensus logo generated based on the sequences of 232 proteins of the Pet20 protein family (InterPro: IPR014804; Pfam: PF08692). Stretches of conserved N_Pet20_- and C_Pet20_-motifs are highlighted by a blue rectangle. Residues are colored based on their physico-chemical properties. (**B**) N_Pet20_- (left) and C_Pet20_-motif (right) of Mrx6 (color intensity corresponds to level of conservation) and the IPR014804/PF08692 consensus (bottom). Residues predicted for hydrogen bond formation (S5C+S7E) or hydrophobic interactions (S7F) to the Pim1 NTD are indicated with black and salmon triangles, respectively. (**C-F**) AlphaFold 3 multimer prediction of 6x Pim1 NTD + ATPase (181-770 with deleted disordered region 285-365), 3x Mrx6 (107-524) and 3x Mam33 (47-266) best ranked model (ipTM + pTM confidence of ~0.46). (**C**) Ribbon model of hexameric Pim1 (181-770 Δ285-365) with separately colored NTD (181-528, gray) and ATPase domains (529-770, blue). Acute-(orange-red NTD, blue ATPase) or obtuse-angled (orange NTD, gray ATPase) protomers alternate. (**D**) Ribbon model of predicted structure consisting of 6x Pim1 NTD + ATPase, 3x Mrx6 (dark green with lime Pet20 motifs) and 3x Mam33 (purple). (**E**) As in D, but Mam33 and Mrx6 C-terminus (271-524) are hidden. (**F**) Detail of (E) showing only one obtuse-angled (low opacity orange, chain A) and the adjacent acute-angled (low opacity orange-red, chain B) Pim1 NTD together with the Mrx6 N-terminus (107-270). Residues involved in hydrogen bonds are highlighted. (**G**) qPCR on mtDNA encoded *COX1* and nuclear reference *TAF10* of indicated strains grown in YPD at 30°C. n = 4, indicated Mrx6 mutant strains expressing either WT Pim1 (filled circles) or Pim1-Spot (empty circles) were analyzed in two biological replicates each, data represent mean ± SD. (**H**) Top panel: Immunoprecipitation pf Pim1-Spot from from native cell lysates of indicated strains grown in YPG at 30°C. Immunoblots were probed with Pim1, Mrx6, Mam33 and Abf2 (control). 50% of eluted fractions and 1% input and flow-through (FT) of the WT Pim1-Spot controls were loaded (for all 1% input and flow-through (FT) controls see6 S7G). Bottom panel: Immunoblots of denatured total lysates of identical strains cultivated in YPG at 30°C are shown (enhanced detection compared to S7G). C.B.B.: colloidal Coomassie stained high molecular weight proteins that retained in the gel after transfer.

In summary, our findings suggest that up to three Mrx6 proteins may associate with a Mam33 trimer via their C-terminal domains and that this interaction is required for Mrx6 stability. Furthermore, our phenotypic analysis reveals a mutual dependency between Mrx6 and Mam33 in regulating WT-like mtDNA levels.

### A bipartite-motif in the Pet20 domain mediates interaction between Mrx6 and Pim1

Next, we examined how Pim1 interacts with Mrx6. We have previously shown that Pim1 co-purifies with any of the three proteins containing the Pet20 domain (Göke et al., 2020), implying that the Pet20 domain may facilitate these interactions. An alignment of 232 proteins annotated for the Pet20 domain family revealed a bipartite nature of the Pet20 domain, which includes two conserved motifs - one located at the N- (~29 AA) and another at the C-terminus (~33 AA) of the Pet20 domain (hereafter N_Pet20_- and C_Pet20_-motif) (Figures 3A+B and Supplementary Figures S5A+B). The N_Pet20_- and C_Pet20_-motifs are separated by a variable region comprising 68 residues in Mrx6, 27 in Pet20 and 63 in Sue1 (Supplementary Figures S1A-C). Similar to the N-terminal segment of Mrx6, which emcompasses the Pet20 domain, AlphaFold predictions revealed low confidence scores for the entire Pet20 and Sue1 structures (Figures 2C+D and data not shown). To assess whether these unstructured regions in the Pet20 domains were indicative of a lack of structural templates for AlphaFold predictions or were intrinsically disordered, we determined disorder propensities using flDPnn predictions (Hu et al., 2021). Indeed, high flDPnn disorder propensities were calculated in regions outside the bipartite Pet20 motif (Supplementary Figures S1A-C).

We used structural predictions to reveal potential interactions between Mrx6, Pet20 or Sue1 and Pim1. Pim1 comprises an N-terminal (NTD) (residues 181-528), ATPase (residues 529-838) and protease (residues 893-1133) domain (Supplementary Figure S1D). Complexity and character limitations in earlier AlphaFold versions precluded prediction of multimer interactions of hexameric Pim1 (Yang et al., 2022). Predictions of interactions between Mrx6 and monomeric Pim1 each lacking their MTSs failed to yield high-confidence structural models. Consequently, we broke down Pim1 into its functional domains to explore possible interactions with proteins containing the Pet20 domain. Although no interactions were identified between Mrx6 and the ATPase or protease domains of Pim1, AlphaFold predicted interactions between a well-structured region of the Pim1 NTD and Mrx6, Pet20 or Sue1. These interactions engaged either the N_Pet20_- or C_Pet20_-motifs (Supplementary Figures S4A+B).

Pim1’s NTD comprises a globular domain connected to the ATPase domain by a long helix (LH) (Li et al., 2021). Both features are also observed in AlphaFold predictions. Multiple hydrogen bonds were predicted between Pim1 and the Pet20 domain proteins (Figure 3B and Supplementary Figures S5A-C and S6A-D). A distinctive feature involved in the interaction with the bipartite Pet20 motif is a negatively charged cavity of the Pim1 NTD, located at the kink between the LH and the globular domain, which contains a glutamate at position 479.

In predictions showing the binding of the N_Pet20_-motif to Pim1, the cavity is occupied by the positively charged R135, R127, or R82 of Mrx6, Pet20, and Sue1, respectively, where hydrogen bonds are observed between the terminal amino group within the guanidinium group of these arginines and the hydroxyl oxygen of the E479 carboxylate side chain of Pim1. In all predictions for Mrx6, but only in some for Pet20 and Sue1, an additional stabilizing bond is formed between the imine-like nitrogen within the guanidinium group of the arginine side chain and the carbonyl oxygen of the peptide bond backbone at V448. Another interaction site between Pim1 and the N_Pet20_-motif involves hydrogen bonds between both the side chain and peptide backbone of Pim1(T253) and Mrx6(T117), Pet20(S108 and T109), or Sue1(T64). The residues of the N_Pet20_-motif involved in these two interaction surfaces are connected by an alpha-helix comprising 11 amino acids: N122-A132 (Mrx6), Q114-S124 (Pet20), or A69-S79 (Sue1), none of which appear to exhibit binding in predictions with a single Pim1 NTD.

In predictions where the C_Pet20_-motif interacts with Pim1, the E479 cavity is similarly important. Here, the terminal hydroxyl oxygen of Pim1(E479) accepts protons from the hydroxyl hydrogen of the serine and threonine side chains in Mrx6(S215 and S217), Pet20(S166 and T168), or Sue1(S166). Additionally, the carbonyl oxygen of the peptide bond backbone at V448 (Pim1) acts as a hydrogen bond acceptor, forming interactions with the amide hydrogen of the peptide bond in Mrx6(I216), Pet20(A167), or Sue1(S166).

Interestingly, AlphaFold similarly predicted an interaction between the Mrx6 C_Pet20_-motif and the NTD of the human Lon protease LONP1, while no interactions involving the N_Pet20_-motif were predicted (Supplementary Figures S4A+B and S6E+F). We assessed the conservation of interacting Pim1/LONP1 residues by generating a probabilistic Hidden Markov model consensus logo (Wheeler et al., 2014) for 34 eukaryotic Lon protease homologues, spanning all major eukaryotic supergroups (Supplementary Figures S5D). Particularly Pim1(E479) (E369 in human) is present in a majority of species. Further analysis through progressive-iterative alignment of the 34 sequences (Edgar, 2004) revealed conservation of most Pim1 residues engaged in Pet20 motif binding across Fungi, *C. elegans* and *H. sapiens* (Supplementary Figure S5E).

Recent advancements via AlphaFold 3 (Abramson et al., 2024) allowed the prediction of an oligomeric assembly of the Pim1 hexamer, truncated to its NTD and ATPase domains, along with three Mrx6 and Mam33 molecules (Figures 3C-E and Supplementary Figure S7A-D). The predicted hexameric structure of Pim1’s NTD resembled previously solved structures for Lon’s NTDs in *Thermus thermophilus* (Coscia & Löwe, 2021), *Meiothermus taiwanensis* (Li et al., 2021) and *Homo sapiens* (Mohammed et al., 2022; Kunová et al., 2024). A characteristic of the hexameric NTD assembly is a pseudo-threefold arrangement, wherein alternating subunits adopt distinct conformations. At the core, an interlocked helix triangle is formed by three subunits, with LHs extending the first helix of the ATPase domain at an obtuse angle (chains A, C, E), spanning over the core. The LHs of the remaining three protomers are oriented in an acute angle towards the first ATPase domain helix, facing outwards without forming inter-contacts (chains B, D, F). Remarkably, the three Mrx6 molecules symmetrically positioned on top of the Pim1 hexamer, with Pet20 domains extending into a cavity created by the NTD’s of Pim1. Each Mrx6 promoters’s N_Pet20_-motif engages an acute-angled Pim1 NTD while the C_Pet20_-motif engages an adjacent obtuse-angled NTD. These interactions are mediated by the same hydrogen bonds inferred from only pairwise predictions between Pim1’s NTD and either Mrx6, Pet20 or Sue1 (Figure 3F and Supplementary Figures S7E).

Given the presence of multiple conserved hydrophobic residues within the bipartite Pet20 motif, we examined the contributions of hydrophobic side chains to the interaction between Pim1 and the Pet20 motifs. Hydrophobic regions within Mrx6’s N_Pet20_- and C_Pet20_-motifs associate with conserved hydrophobic sites of Pim1’s NTD, which are largely conserved among eukaryotic Lon proteases (Figure 3A and Supplementary Figures S5D+S7F). In particular, the alpha-helix of Mrx6’s N_Pet20_-motif, not crucial for binding in single Pim1 NTD predictions, contains hydrophobic residues Mrx6(L130 and F131) and serves to bridge the acute-angled Pim1 NTD (site of N_Pet20_-motif interaction) to the adjacent obtuse-angled NTD, involving Pim1’s residues V394-F406, I434, F435 and L441 (see Supplementary Figure S7 for further structural information).

The oligomeric prediction revealed no interactions between Pim1 and the Mrx6 C-extension, which binds exclusively to Mam33. Analogous to a closing lid, the trimeric Mam33 is placed on top of the assembly with its highly negatively charged site facing the positively charged Mrx6 C-terminus.

To confirm the predicted interaction sites between Mrx6 and Pim1 *in vivo*, we generated yeast strains containing mutant variants of *MRX6* at its endogenous locus. Specifically, we removed Mrx6’s Pet20 domain (Δ112-240) or replaced it with the corresponding Pet20 domain of Pet20 (Δ112-240::Pet20(104-190)).

Additionally, we generated strains with point mutations in either the N_Pet20_- or C_Pet20_-terminal regions of the bipartite motif at locations predicted to form hydrogen bonds with the Pim1 NTD. In the N_Pet20_-motif, the stabilizing hydrogen bonding between Mrx6(T117) and Pim1(T253 or S254) was prevented by Mrx6^T117V^. The predicted interaction between Mrx6(R135) in the N_Pet20_-motif and Pim1(E479 and V448) was altered by mutating Mrx6’s arginine to alanine or glutamate. Binding of Mrx6’s C_Pet20_-motif to the Pim1(E479) was perturbed by mutating Mrx6(S217) alone or in combination with Mrx6(S215) to alanine. A mutant lacking the C-terminal domain of Mrx6 (Mrx6^ΔC-term.^) was also included in these analyses.

First, we assessed the functional impact of Mrx6 mutants on mtDNA CN level in WT and Pim1-Spot backgrounds. While cells expressing Mrx6^T117V^, Mrx6^R135A^ and Mrx6^S217A^ showed normal mtDNA levels, cells expressing Mrx6^Δ112-240^, Mrx6^Δ112-240::Pet20(104-190)^, Mrx6^R135E^or Mrx6^S215A/S217A^ exhibited increased mtDNA CN comparable to Δ*mrx6* cells, indicating a critical role of Mrx6’s Pet20 domain and the respective residues in facilitating Mrx6 function (Figure 3G).

Next, we performed immunoprecipitations of Pim1-Spot to investigate whether the observed increase in mtDNA CN was associated with a loss of Pim1-Mrx6 interactions. Mrx6 point mutants were stably expressed and detected in a WT-like range in total cell lysates (Figure 3H and Supplementary Figure S7G). Mrx6^Δ112-240^ and Mrx6^Δ112-240::Pet20(104-190)^ were detected at expected sizes. Reduced levels of these mutants cannot be quantitatively compared to WT as only epitopes within 34-111 are present in these variants, while the antibody was generated against residues 34-210. Across all samples, input and eluted amounts of Pim1-Spot were similar. Remarkably, we observed substantial differences in the amounts of immunoprecipitated Mrx6 and Mam33. In line with elevated mtDNA CN levels, we detected no interactions of Mrx6 or Mam33 with Pim1-Spot in strains expressing Mrx6^Δ112-240^ or Mrx6^Δ112-240::Pet20(104-190))^ and a severely weakened interaction between Pim1-Spot with Mrx6^R135E^. Binding of Mrx6 to Pim1-Spot was also reduced in Mrx6^T117V^, Mrx6^R135A^ and Mrx6^S217A^ mutants. Since these mutants exhibit largely unaltered mtDNA CN, we conclude that compromised yet not completely abrogated affinity between Mrx6 and Pim1 is sufficient to maintain WT-like mtDNA levels. We hypothesize that hydrophobic interactions, such as those involving nearby Mrx6(Y134) or Mrx6(I216), are sufficient to sustain the Pim1-Mrx6 interaction in the absence of hydrogen bonds from Mrx6(R135) and Mrx6(S217), respectively. In accordance with our observation that the association of Mam33 with Pim1 is dependent on Mrx6, bound Mam33 levels decreased proportionally along with Mrx6. It is notable that Mrx6^R135A^ and Mrx6^S215A/S217A^ exhibit comparable binding deficits to Pim1. However, while Mrx6^S215A/S217A^ results in strongly increased mtDNA levels, Mrx6^R135A^ does not, indicating that the roles of Mrx6(S215 and S217) may extend beyond merely facilitating interaction between Mrx6 and Pim1.

Taken together, a bipartite motif within the Pet20 domain of Mrx6 facilitates interaction with the NTD of Pim1. Central to this interaction is a negatively charged cavity formed by E479 in Pim1. Our *in vivo* results highlight the crucial role of Mrx6(R135) in the N_Pet20_-motif and Mrx6(S215, S217) in the C_Pet20_-motif in facilitating Pim1 binding and regulating mtDNA copy number.

### Absence of Mrx6 leads to a post-transcriptional increase of Rpo41 and Cim1

Our data suggest a model in which the binding of Mrx6, along with Mam33, to Pim1 alters recruitment and/or degradation of factors required for mtDNA CN maintenance. To identify proteins that may accumulate in the absence of Mrx6, we isolated mitochondria-enriched fractions from WT and Δ*mrx6* cells and conducted a comparative proteomic analysis. Comparison of the obtained dataset with a high-confidence mitochondrial proteome (Morgenstern et al., 2017) and a SILAC dataset profiling Δ*mam33* and WT cells (Hillman & Henry, 2019), revealed good coverage of the mitochondrial proteome as well as proteins annotated for maintenance of mtDNA and the nucleoid (Cho et al., 1998; Meeusen et al., 2000; Kaufman et al., 2000; Sato et al., 2002; Sakasegawa et al., 2003; Sato & Miyakawa, 2004; Chen et al., 2005) (Figure 4A and Supplementary Figures S8E-H).

**Figure 4.**
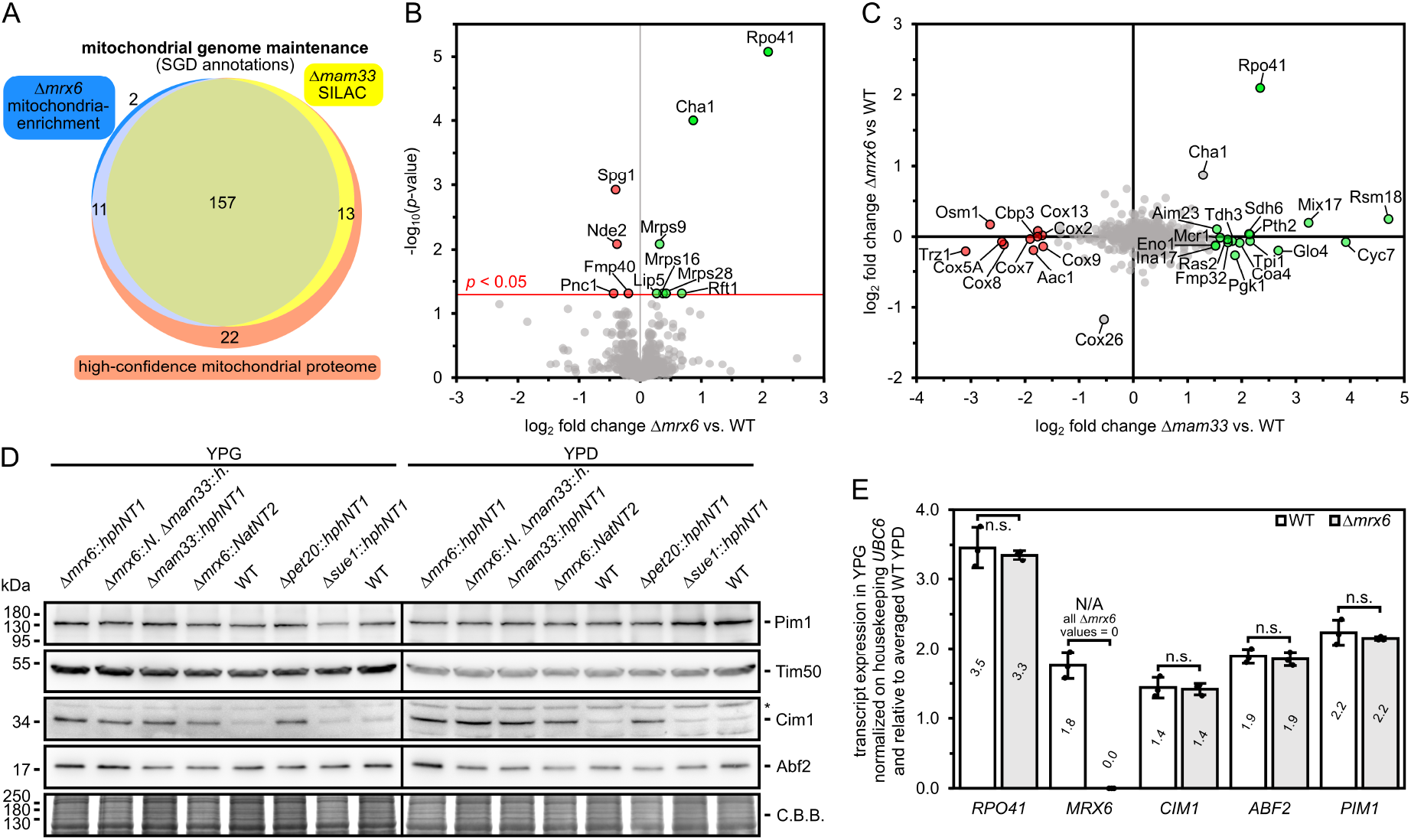
Absence of Mrx6 leads to a post-transcriptional increase of Rpo41 and Cim1. (**A**) Comparison of proteins annotated for mitochondrial genome maintenance (http://www.yeastgenome.org, state February 2024) detected in the proteome of mitochondrial enriched fractions from Δ*mrx6* and WT cells (this study), a SILAC dataset from Δ*mam33* and WT cells filtered for mitochondrial proteins (Hillman & Henry, 2019) and a high-confidence mitochondrial proteome (Morgenstern et al., 2017). (**B**) Volcano plot of differentially expressed proteins identified by LC-MS/MS analysis in mitochondrial enriched fractions of WT and mrx6 cells cultivated in YPG at 30°C. Normalization based on label-free quantification (LFQ). n = 2 strain replicates, two technical replicates each. (**C**) Comparison of proteome alterations present in the Δ*mrx6* vs. WT LFQ LC-MS/MS derived data as well as a Δ*mam33* vs. WT SILAC dataset filtered for mitochondrial proteins (Hillman & Henry, 2019). Fold-changes of at least 2.8 in at least one of the two experiments are highlighted in green or red. (**D**) Immunoblots of Pim1, Tim50 (loading control), Cim1, and Abf2 from lysates of indicated cells grown in YPG or YPD at 30°C. * highlights unspecific cross-reaction of the Cim1 antibody. C.B.B.: colloidal Coomassie stained high molecular weight proteins that retained in the gel after transfer. n = 1. (**E**) RT-qPCR transcript comparison of indicated genes between WT and Δ*mrx6* with RNA extracted from cells cultivated in YPD or YPG medium at 30°C. Normalization to housekeeping gene *UBC6* and relative to the averaged WT abundance in YPD (see S8A+B for only YPD and YPG). n = 3 biological replicates, data represent mean *±* SD.

While most of the 931 identified proteins were not altered between WT and Δ*mrx6* samples, two nucleoid proteins, the mitochondrial RNA polymerase Rpo41 (Greenleaf et al., 1986) and the catabolic serine and threonine deaminase Cha1 (Petersen et al., 1988), were markedly increased ~4.3-fold and ~1.8-fold, respectively, in the absence of *MRX6* (Figure 4B). Given our finding that both the absence of either *MRX6* or *MAM33* increases mtDNA CN levels, we compared our Δ*mrx6* dataset with a previously published dataset on protein changes associated with deletion of *MAM33*, to identify common changes. Indeed, the levels of Rpo41 and Cha1 were similarly increased by ~5.1-fold and ~2.5-fold, respectively, in Δ*mam33* cells (Figure 4C). The protein Cox26 was decreased in Δ*mam33* and Δ*mrx6* cells, but the change was not significant comparing Δ*mrx6* to WT. The abundance of numerous proteins was exclusively altered in Δ*mam33* cells, suggesting that Mam33 has additional functions beyond its role in the Mrx6-Pim1 complex. This idea is supported by stronger growth defects of Δ*mam33* compared to Δ*mrx6* cells and existing literature proposing roles of Mam33 as a *COX1* translation activator (Roloff & Henry, 2015) or a chaperone-like protein facilitating mitochondrial ribosome assembly (Hillman & Henry, 2019). Notably, Cha1, whose steady-state level was elevated in Δ*mrx6* and Δ*mam33* cells, was also detected in our Pim1-Spot pull-down experiments. In these experiments, a minor yet statistically significant interaction with Pim1 was observed in the absence of Mrx6.

We further focused on Rpo41, which showed the strongest steady state protein level increase in Δ*mrx6* and Δ*mam33* cells. The mitochondrial RNA polymerase (RNAP) is homologous to the T3/T7 RNAPs bacteriophages (Masters et al., 1987) and besides its function in mtDNA transcription (Matsunaga & Jaehning, 2004; Deshpande & Patel, 2012; 2014; Sultana et al., 2017) its role in priming mtDNA synthesis by the mtDNA polymerase Mip1 at origins of replication and promoter sequences has been reported (Sanchez-Sandoval et al., 2015; Ramachandran et al., 2016). We quantified Rpo41 protein levels in lysates from WT and Δ*mrx6* cells grown in in YPG (medium used for mitochondria for LC-MS/MS) and YPD by immunoblotting. Rpo41 levels were increased ~2.3-fold (YPG) and ~1.4-fold (YPD) compared to the ~4.3-fold increase observed by LC-MS/MS analysis of YPG mitochondria (Supplementary Figure S8C). This discrepancy may arise from dissimilar linearities in immunoblot and LC-MS/MS analysis or could be attributed to ongoing degradation of Rpo41 during mitochondrial isolation in WT samples, where Mrx6 is present.

Consistent with our LC-MS/MS findings, immunoblots reaffirmed that the mitochondrial proteins Mam33, Pim1, Cor2, Atp2, Cox2, Tim23 and Cox2 remained unchanged in Δ*mrx6* cells. We further assessed the levels of Rpo41, Mrx6 and Mam33 in cells lacking the other Pet20 domain proteins, Pet20 or Sue1, through immunoblotting. The absence of Sue1 did not lead to any changes in the levels of the tested proteins (Supplementary Figure 8D). However, strains devoid of Pet20 exhibited an increased abundance of Rpo41, suggesting that Pet20 also affects Rpo41 degradation.

Despite comprehensive coverage of the mitochondrial proteome and mtDNA maintenance factors, the LC-MS/MS dataset lacks some factors implicated in the maintenance or expression of mtDNA such as Cim1, Mip1, Hmi1, Pif1 and Exo5 and mitochondrial proteins annotated for DNA-binding (Supplementary Figures S8H+I). Therefore, we can not make any statements about the abundance of these proteins in the presence of absence of Mrx6 based on our mass spectrometry results.

We recently demonstrated that Cim1, an HMG-box protein, is essential for mtDNA CN regulation, and its absence results in elevated mtDNA levels (Schrott & Osman, 2023). We had found that Cim1 is maintained at low levels in WT cells but accumulates when Pim1 is depleted. Hence, we tested if Cim1 levels might be affected in cells lacking Mam33, Mrx6 or Pet20. Indeed, Cim1 accumulated in Δ*mrx6*, Δ*mam33* and Δ*pet20* cells, suggesting that Cim1 levels are regulated through degradation by the concerted action of Pim1, Mrx6, Mam33 and Pet20 (Figure 4D).

To rule out transcriptional alterations due to secondary effects triggered by the absence of Mrx6, we isolated RNA from WT and Δ*mrx6* cells and performed reverse transcriptase qPCR (RT-qPCR) to quantify transcript levels of *RPO41* as well as *MRX6, CIM1, ABF2* or *PIM1*. All these factors involved in mtDNA maintenance were generally upregulated in respiratory medium (YPG) compared to fermentable medium (YPD), but we did not observe significant differences between WT and Δ*mrx6* for these transcripts (Figure 4E and Supplementary Figures S8A+B).

In summary, absence of any of the Pim1 complex partners Mrx6, Mam33 or Pet20 results in increased protein abundance of the mitochondrial RNA polymerase Rpo41 and the HMG-box protein Cim1, known factors involved in mtDNA maintenance, without changes of their transcript levels. Additionally, our results suggest that Cha1 is a Pim1 substrate that is stabilized in the absence of Mrx6 or Mam33.

### Absence of Mrx6 or Pim1 increases the lifetime of Rpo41

Mrx6 interacts with Pim1’s N-terminal domain, which is proposed to mediate substrate recognition (Coscia & Löwe, 2021), implying that Mrx6 might facilitate the degradation of Rpo41 and Cim1 by Pim1, as suggested by their increased levels in Δ*mrx6* cells. We performed cycloheximide-chase experiments to test this idea. Due to the low abundance of Cim1 in WT, reliable detection by immunoblotting is challenging, which also hinders the assessment of further decreases during chase experiments. Therefore, we assessed only the stability of Rpo41 in WT and Δ*mrx6* cells during different chase periods following inhibition of cytosolic translation with cycloheximide. Within four hours post-cycloheximide addition, the abundance of Rpo41 and, remarkably also Mrx6 rapidly decreased, in contrast to Mam33, whose levels stayed constant. Higher levels of Rpo41 were consistently detected in Δ*mrx6* cells, yet no difference in its decay was noted compared to WT (Figures 5A-D).

**Figure 5.**
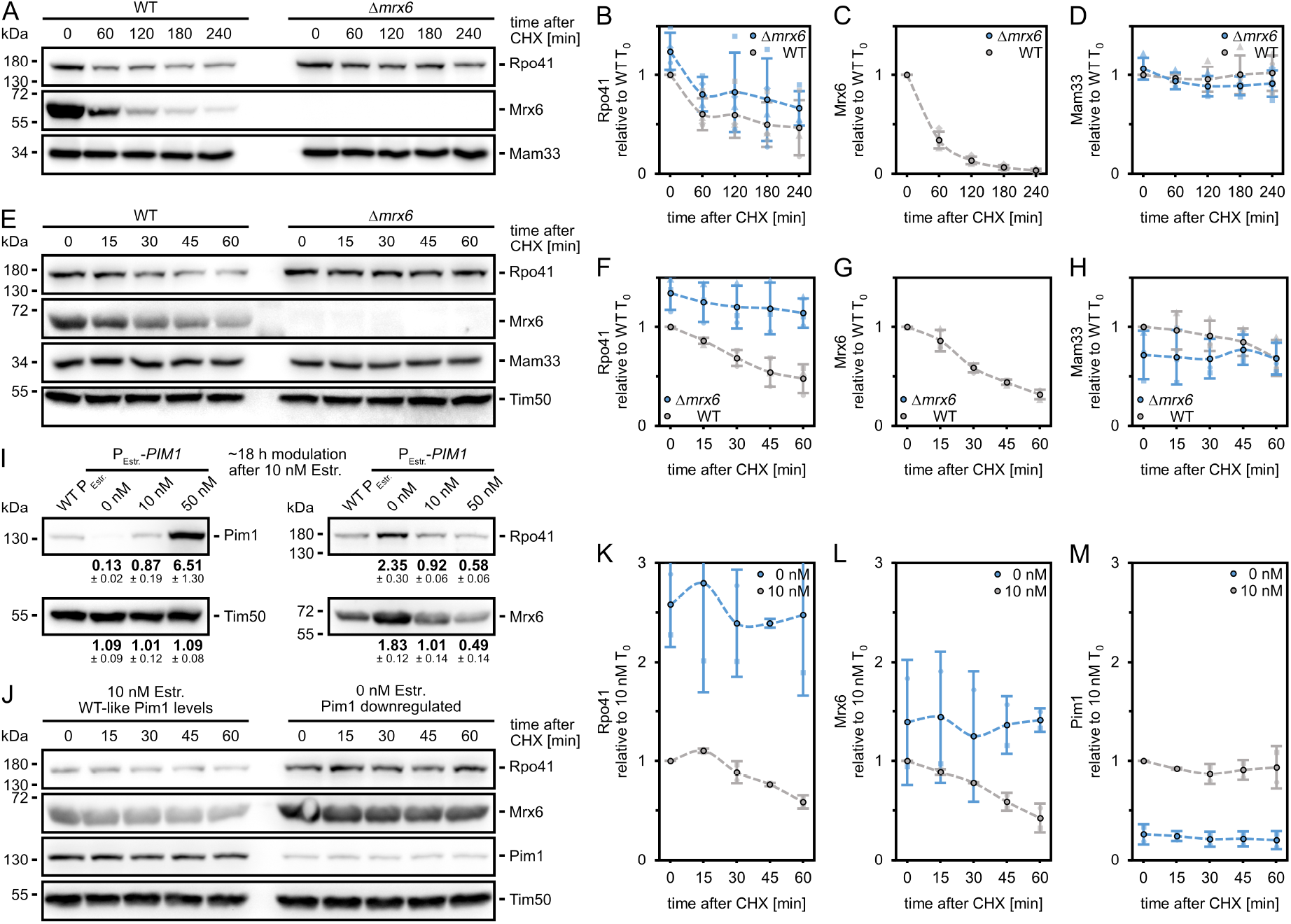
Absence of either Mrx6 or Pim1 increases the lifetime of Rpo41. (**A-H**) Immunoblots and quantifications of Rpo41, Mrx6, Mam33 and Tim50 (additional loading control in E-H, see S9A for quantification) from lysates of WT and Δ*mrx6* cells cultivated in log phase in YPG at 30°C and chased for four hours (A-D) or one hour (E-H) after addition of cycloheximide (CHX). n = 4 (A-D) and n = 3 (E-H), data represent mean (dots with black outline) ± SD. (**I**) Immunoblots and quantifications of Pim1, Rpo41, Mrx6 and Tim50 (loading control) from lysates of cells expressing Pim1 from the endogenous promoter and containing an empty P_Estr._ promoter construct and cells containing a P_Estr._-*PIM1* construct cultivated in log phase in YPG supplemented for ~18 h with 0, 10 or 50 nM estradiol at 30°C. n = 3 biological replicates, data represent mean *±* SD. (**J-M**) Immunoblots and quantifications of Rpo41, Mrx6, Pim1 and Tim50 (loading control, see S9B for quantification) from lysates of P_Estr._-*PIM1* cells cultivated in log phase in YPG at 30°C and chased for one hour after addition of CHX. WT-like Pim1 levels were sustained with 10 nM estradiol, whereas prior withdrawal of estradiol for ~18 h resulted in downregulation. n = 2, data represent mean (dots with black outline) *±* SD.

However, when we focused on dynamics within the first hour after addition of cycloheximide in 15-minute intervals, Rpo41 exhibited a slower degradation rate in Δ*mrx6* compared to WT cells. In contrast, the levels of Mam33 and Tim50 remained stable (Figures 5E-H and Supplementary Figure S9A). We used an estradiol-inducible promoter integrated upstream of the endogenous *PIM1* ORF to examine whether direct modulation of Pim1 levels similarly affects Rpo41 stability (Schrott & Osman, 2023). In respiratory medium, the addition of 10 nM of estradiol resulted in WT-like Pim1 levels. Omitting estradiol for ~18 hours in log phase cultures overnight caused a ~7.7-fold decrease in Pim1 levels and a ~2.4-fold or ~1.8-fold increase in Rpo41 and Mrx6, respectively, while Tim50 remained unchanged (Figure 5I).

A ~6.5-fold increase in Pim1 upon 50 nM estradiol vice versa decreased Rpo41 levels ~1.7-fold and Mrx6 ~2-fold. When we combined the downregulation of Pim1 with an one-hour cycloheximide chase, Rpo41 and Mrx6 protein levels were stabilized (Figures 5J-M and Supplementary Figure S9B).

Our results strongly suggest that Pim1 and Mrx6 participate in the degradation of Rpo41. For this process to occur, Rpo41 must at least transiently associate with the Pim1-Mrx6 complex. Supporting this notion, recent immunoprecipitation experiments using overexpressed catalytically inactive Pim1^S1015A^-HA alongside endogenous Pim1 demonstrated co-purification of Rpo41 with Pim1 (Bertgen et al., 2024). However, our LC-MS/MS analysis of proteins co-purifying with catalytically active Pim1-Spot did not identify Rpo41 as a significant hit (Figure 1D). Only three unique Rpo41 peptides were detected in two out of three replicates from Δ*mrx6* samples, while no peptides were found when Mrx6 was present. Considering the findings of Bertgen et al. alongside our own results, it appears that the interaction between Rpo41 and Pim1 is stabilized when Pim1’s catalytic activity is impaired, suggesting that efficient substrate processing may prevent prolonged association.

To confirm this notion and to examine the role of Mrx6 in association of Rpo41 with catalytically impaired Pim1^S1015A^, we performed immunoprecipitation experiments in WT or Δ*mrx6* cells overexpressing HA-tagged Pim1^S1015A^ in addition to endogenous Pim1. Co-purification of Rpo41 with Pim1^S1015A^ was observed by Western Blot (Supplementary Figure S9C). Interestingly, the Pim1^S1015A^-Rpo41 association was independent of Mrx6, since similar amounts of Rpo41 co-purified with Pim1 in Δ*mrx6* cells. Like in previous experiments (Figure 1A-D), co-purification of Mam33 with Pim1 was dependent on Mrx6.

We next investigated whether Mrx6 levels are rate-limiting for Rpo41 degradation and the regulation of mtDNA copy number. To address this, we utilized a system enabling estradiol-dependent expression of *MRX6* from its native locus. In respiratory medium lacking estradiol, cells retained approximately 50% of WT Mrx6 levels, likely due to basal leakiness of the estradiol-inducible promoter. To assess the impact of increased Mrx6 levels, we induced its expression with 20 nM and 100 nM estradiol for two hours, resulting in approximately 6-fold and 12-fold increases in Mrx6 levels, respectively (Supplementary Figure S9D). Despite substantial overexpression, neither condition significantly affected mtDNA copy number or altered the abundance of Rpo41 or Mam33, suggesting that Mrx6 levels are not a limiting factor for Rpo41 degradation.

In summary, our findings suggest that Rpo41 and Mrx6 are degraded by the Pim1 protease. Although Rpo41 still binds to Pim1 in the absence of Mrx6, its stability increases when Mrx6 is missing, suggesting that Mrx6 supports Pim1-mediated degradation of Rpo41.

### Absence of Mrx6 suppresses Cim1 function

We sought to better understand how Mrx6 affects mtDNA levels. We have determined that absence of Mrx6 and downregulation of Pim1 increase Rpo41 abundance and delayed its degradation. Increased Rpo41 amounts may elevate the frequency of priming events necessary for mtDNA replication, resulting in increased mtDNA CN in Δ*mrx6* cells. To disentangle elevated Rpo41 levels from other alterations that absence of Mrx6 might impart, we engineered strains where the expression of *RPO41* can be modified via estradiol. Supplementing YPG medium with 10 nM estradiol resulted in expression levels comparable to WT, whereas concentrations below or exceeding 10 nM led to downregulation or upregulation of Rpo41, respectively (Supplementary Figure S9E). To exclude cellular adaptation processes, we first acutely modulated *RPO41* expression for only six hours. While Mrx6 and Mam33 remained largely unchanged, Rpo41 levels increased ~2-fold or ~4-fold in the presence of 15 and 30 nM estradiol, respectively, and its abundance halved in the absence of estradiol (Figure 6A and Supplementary Figure S9F). Rpo41 upregulation did not significantly affect mtDNA CN, but downregulation reduced mtDNA CN. To mimic steady state elevated Rpo41 of Δ*mrx6* cells, we doubled Rpo41 levels with 15 nM estradiol over ~20 hours and observed a slight increase (~1.16-fold) in mtDNA CN (Figure 6B). However, this increase was indistinguishable from control strains that expressed the estradiol-receptor-fused transcription factor, but did not contain responsive elements in the *RPO41* promoter, and therefore expressed WT Rpo41 levels, irrespective of the presence of Estradiol. These results indicate that Rpo41 upregulation alone does not increase mtDNA levels.

**Figure 6.**
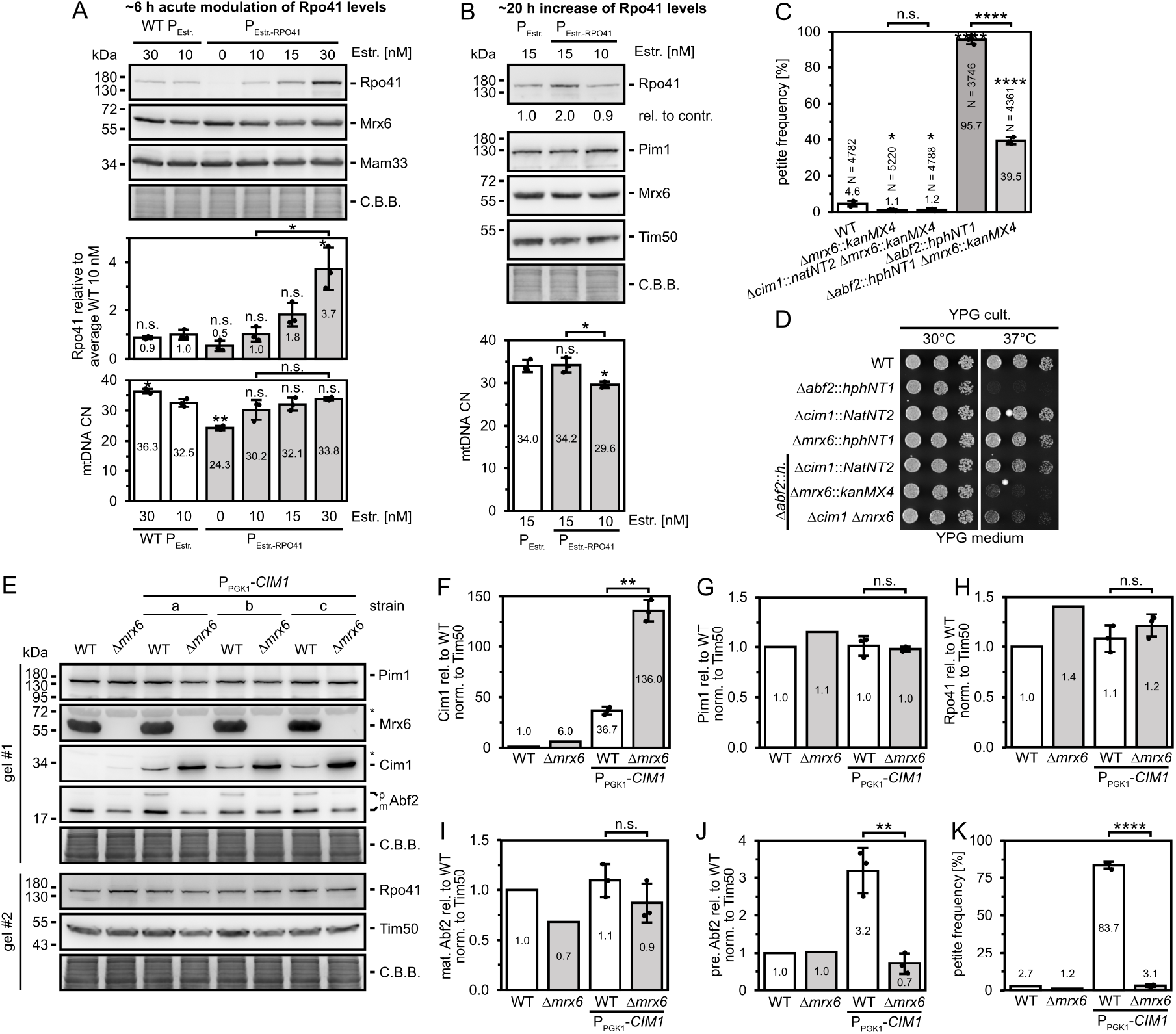
Absence of Mrx6 suppresses Cim1 function. (**A+B**) Immunoblots of Rpo41 (with quantifications), Mrx6 and Mam33 (loading control) of lysates from P_Estr._-*RPO41* cells grown in YPG at 30°C for ~6 h (A) or for ~20 h (B) with indicated estradiol concentrations starting from 10 nM estradiol (WT-like Rpo41 levels). Cells with WT *RPO41* promoter lacking LexA binding sites but expressing the LexA-(estradiol-LBD)-B42 activator served as control (WT P_Estr._). C.B.B.: colloidal Coomassie stained high molecular weight proteins that retained in the gel after transfer. Rpo41 signals were quantified relative to the averaged signal of WT P_Estr._ cells cultivated with 10 nM (A) or 15 nM (B) estradiol. mtDNA CN was determined by qPCR on mtDNA encoded *COX1* and nuclear reference *TAF10*. n = 3, data represent mean ± SD. (see S9G for quantifications of Mrx6 and Mam33 signals in A) (**C**) Petite frequency of indicated strains grown in YPD at 30°C. Experiment was performed as published and data for WT and Δ*abf2* are identical to (Schrott & Osman, 2023). n = 3 biological replicates, data represent mean ± SD. (**D**) Drop dilution growth analysis at 30°C or 37°C of indicated strains pre-grown in YPG at 30°C. Images were taken after 48 h growth on YPG (for YPD see S9G). Only dilution steps 2-4 are shown. (**E-J**) Immunoblots (and quantifications) of Mrx6, Cim1 (F), Pim1 (G), Rpo41 (H), Abf2 including premature (p) and mature (m) forms (I+J) and Tim50 (loading control) of lysates from indicated cells grown in YPD at 30°C. For P_PGK1_-*CIM1* overexpression in either WT or Δ*mrx6* backgrounds n = 3 independent strains a, b and c were analyzed. * highlights unspecific cross-reaction of Cim1 and Mrx6 antibodies. C.B.B.: colloidal Coomassie stained high molecular weight proteins that retained in the gel after transfer. (**K**) Petite frequency of indicated strains grown in YPD at 30°C. n = 3 (strains a, b and c) datapoints of P_PGK1_-*CIM1* samples represent two averaged biological replicates each, WT and Δ*mrx6* n = 1, data represent mean *±* SD.

Since elevated Rpo41 levels do not appear to be mainly responsible for elevated mtDNA levels in Δ*mrx6*, we investigated the interplay between Mrx6 and Cim1, whose levels also rise in cells lacking Mrx6, Mam33, Pet20, or upon Pim1 downregulation. The increase of Cim1 levels was unexpected in Δ*mrx6* cells exhibiting increased mtDNA levels, since we have reported previously that higher Cim1 levels result in reduced mtDNA CN and drastically increased petite frequencies (Schrott & Osman, 2023). Cells lacking functional Cim1, by contrast, display elevated mtDNA CNs and lower petite frequencies, similar to deletions of *MRX6* and *MAM33* (Figure 2H) (Schrott & Osman, 2023). Therefore, we hypothesized that the increase of mtDNA CN in cells lacking an intact Pim1-Mrx6 complex is due to impaired Cim1 functions. In line with this idea, petite frequencies in Δ*mrx6*Δ*cim1* double mutants were indistinguishable from Δ*mrx6* (Figure 6C) or Δ*cim1* (Schrott & Osman, 2023) single mutants, pointing to an epistatic relationship.

To further test this idea, we first examined the effect of Mrx6 loss on cells lacking the mtDNA packaging factor *ABF2*. Δ*abf2* cells exhibit a high petite frequency when grown in YPD and lose the ability to respire at 37°C (Diffley & Stillman, 1991; Schrott & Osman, 2023). Both phenotypes can be rescued by deletion of *CIM1*, revealing a detrimental function of Cim1 on mtDNA when Abf2 is absent (Schrott & Osman, 2023). If absence of Mrx6 impairs Cim1 function, we reasoned that deletion of Mrx6 might also rescue phenotypes associated with loss of Abf2. Indeed, deletion of *MRX6* was sufficient to reduce high petite frequencies of Δ*abf2* cells cultivated in YPD and to partially restore respiratory growth of Δ*abf2* cells at 37°C (Figure 6C+D, Supplementary Figure 9G). However, Δ*abf2*Δ*mrx6* cells grew worse than Δ*abf2*Δ*cim1* double mutants. This observation suggests that Cim1 may retain partial functionality in the absence of Mrx6 or that Mrx6 has additional roles that become important when Abf2 is absent — possibilities that are not mutually exclusive. The Δ*abf2*Δ*cim1*Δ*mrx6* triple mutant grew slightly better than the Δ*mrx6*Δ*abf2* double mutant at 37°C on YPG, suggesting that Cim1 remains partially functional in the absence of Mrx6. On the other hand, the Δ*abf2*Δ*cim1*Δ*mrx6* triple mutant grew less well than the Δ*cim1*Δ*abf2*, suggesting that Mrx6 fulfills functions beyond suppression of Cim1. Such a function could relate to Mrx6’s role in affecting Rpo41 stability.

To provide further evidence for a suppressive effect of Mrx6 absence on Cim1 function, we asked whether deletion of *MRX6* would alleviate high petite frequency caused by overexpression of Cim1. Hence, we expressed Cim1 from a *PGK1* promoter at the genomic *CIM1* locus (Schrott & Osman, 2023). In WT cells, Cim1 levels increased by ~37-fold and further accumulated ~136-fold in Δ*mrx6* cells without affecting abundances of Mrx6, Pim1, Rpo41 or the mature form of Abf2 (Figures 6E-I). Surprisingly, although Cim1 levels were even higher in three independent P_PGK1_-*CIM1* overexpression strains generated in the Δ*mrx6* background compared to WT, the accumulation of Abf2 precursors and dramatically increased petite frequencies were entirely rescued in the absence of *MRX6* (Figures 6E, J, K).

Taken together, increasing levels of Rpo41 artificially in WT cells to levels that are observed in Δ*mrx6* cells did not affect mtDNA CN. Surprisingly, our results reveal that the effect of Mrx6 on mtDNA is linked to Cim1, which does not cause inhibitory effects on mtDNA maintenance when Mrx6 is absent.

## DISCUSSION

In this study we provide evidence for a complex consisting of the conserved Lon protease, Pim1, the Pet20 domain containing protein Mrx6 and Mam33. Our data suggest that a Mrx6-Mam33 subcomplex binds to the N-terminal domain of Pim1 and determines Pim1’s activity or specificity towards selective substrates. Absence of Mrx6 or Mam33 as well as point mutations impairing the Pim1-Mrx6 interaction lead to increased mtDNA CN. Furthermore, the mitochondrial RNA polymerase, Rpo41, and the mitochondrial HMG-box protein Cim1 accumulate to higher levels in Δ*mrx6* or Δ*mam33* cells. Strikingly, all components have been linked to mtDNA CN regulation. Hence, our data reveal insight into a complex network governing mtDNA CN.

We have previously hypothesized that Mrx6 regulates substrate degradation by Pim1 to control mtDNA CN (Göke et al., 2020). Accumulation of Rpo41 and Cim1 in the absence of *MRX6* supports this idea. Rpo41 is a mitochondrial RNA polymerase that is also required for priming of mtDNA replication (Ramachandran et al., 2016; Sanchez-Sandoval et al., 2015; Baldacci et al., 1984). Elevated levels of Rpo41 could therefore result in an increase of priming events and elevated mtDNA CN. However, we show that an increase of Rpo41 is insufficient to increase mtDNA levels by its own, hinting towards a multilayered regulatory network. Surprisingly we find that also Cim1 accumulates in the absence of Mrx6, revealing that Cim1 degradation is regulated by Mrx6. Furthermore, Mrx6 also modulates Cim1 function, as deletion of Mrx6 mitigates negative effects on the maintenance of mtDNA associated with Cim1 overexpression. How Mrx6 impacts Cim1 function is a remaining question, but could involve post-translational modifications, altered protein interactions or submitochondrial localization changes. It is possible that Mrx6 modulates the degradation of additional Pim1 substrates within a broader regulatory network. One potential candidate is Cha1, which accumulates in Δ*mrx6* or Δ*mam33* cells (Hillman & Henry, 2019). While deletion of Cha1 did not increase petite frequencies, its acute overexpression raised them by approximately twofold (data not shown), suggesting a potential role in mtDNA maintenance. Such a link is further supported by the observation that Cha1 associates with mtDNA (Chen et al., 2005). How Cha1 is involved in the process and how this may involve Cha1’s function as a Serine and threonine deaminase (Petersen et al., 1988) remains to be determined.

How may the Mrx6-Mam33 subcomplex affect degradation of substrates by Pim1? While deletion of Pim1 results in respiratory deficient yeast cells (Van Dyck et al., 1994; Suzuki et al., 1994), deletion of Mrx6 or Mam33 results in yeast cells capable of respiratory growth. Thus, Mrx6 and Mam33 do not affect all functions of Pim1. This hypothesis is supported by our observation and those of others (Bertgen et al., 2024) that only a subset of Pim1 interacts with Mrx6 or other Pet20 domain containing proteins. It has been proposed that the N-terminal domain of Pim1 is involved in discrimination of substrates, how discrimination is accomplished, however, remains unknown. Parts of the N-terminal domain of Pim1 protomers that are close to its AAA-domain have been suggested to function as a ruler to prevent access and degradation of substrates with insufficiently long exposed C-termini (Li et al., 2021). The binding of Mrx6 to Pim1’s N-terminus may induce conformational changes in Pim1 or its substrates, such as Rpo41 or Cim1, thereby modulating their access to Pim1’s AAA domain and facilitating the initiation of their degradation (Figure 7). Importantly, our data show that recruitment of Rpo41 to Pim1 is independent of Mrx6 but that its degradation is enhanced in the presence of Mrx6. At least in the case of Rpo41, recruitment and regulation of degradation therefore seem to be separable events.

**Figure 7.**
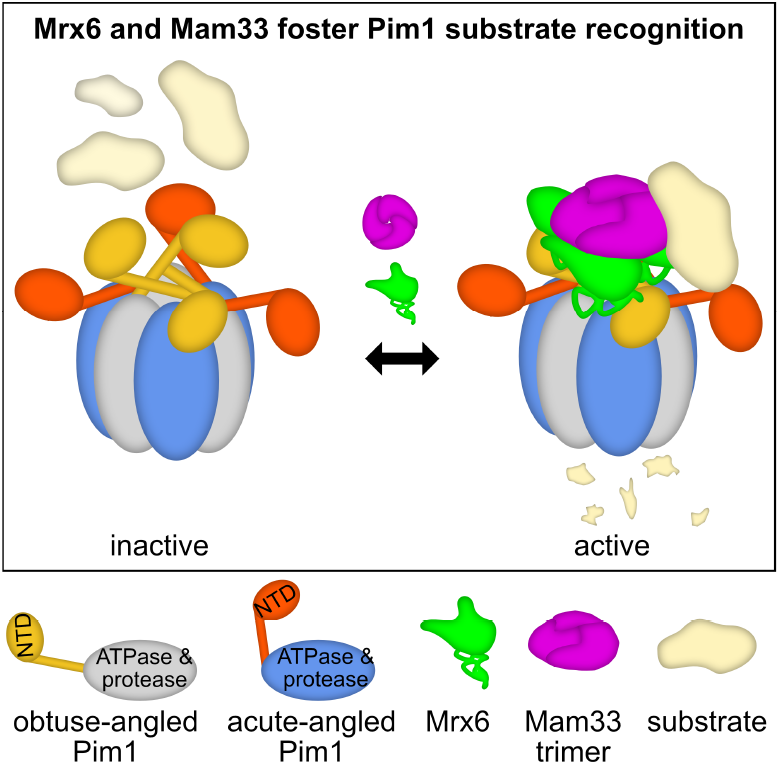
Degradation of Pim1 substrates involved in mtDNA maintenance, mediated by Mrx6 and Mam33.

An interesting aspect of Mrx6 is that it is rapidly turned over in a Pim1-dependent manner. Possibly, Mrx6 is degraded by Pim1 together with other client proteins. The two homologs of Mrx6, Sue1 and Pet20, also possess the bipartite Pet20 motif, which likely facilitates their binding to Pim1. It is an intriguing idea that binding of Mrx6, Pet20 or Sue1 to Pim1 may affect Pim1 or substrates of Pim1 in different ways. Indeed, Sue1 has been linked to altered stability of cytochrome c (Wei & Sherman, 2004). The results presented here and previously (Göke et al., 2020) suggest that Sue1 binds Pim1 independently of Mrx6. Stable association of Pet20 with Pim1 on the other hand requires Mrx6. A mixed complex containing Mrx6 and Pet20 is therefore imaginable. Surprisingly, we show that deletion of Pet20 results in elevated Rpo41 and Cim1 levels, while it does not increase mtDNA levels to the same extent as deletion of *MRX6* (Göke et al., 2020). These results indicate that Mrx6 serves additional roles that cannot be fulfilled by Pet20.

The Lon protease is emerging as a central player in regulation of mtDNA CN. In humans, a large-scale genetic linkage analysis has revealed that LonP mutations are linked to altered mtDNA CN (Gupta et al., 2023). Moreover, LonP has been shown to be important for determining levels of TFAM, which in turn may affect mtDNA CN (Lu et al., 2013). Intriguingly, the Lon proteases in *Caulobacter crescentus* and *Escherichia coli* have similarly been implicated in regulation of DNA replication. In C. *crescentus*, proteotoxic stress results in Lon-mediated degradation of the replication factor DnaA, which thereby prevents DNA replication (Jonas et al., 2013). In *E. coli* on the other hand, Lon-mediated degradation of the replication inhibitor CspD is suggested to coordinate DNA replication with physiological conditions (Langklotz & Narberhaus, 2011).

How is Mrx6 integrated into homeostatic mtDNA CN regulation? While further experiments are needed, we hypothesize that Mam33 could serve important roles in this context. Mam33 and its homologs (p32/HABP1/gC1qR in mammals) are multifunctional proteins that have been implicated in a variety of processes in various cell compartments, including the immunological response, cell cycle regulation, apoptosis or nuclear transcription (Brokstad et al., 2001; Emelyanov et al., 2014; Saha & Datta, 2018). A mitochondrial function of Mam33 across different species is evident from its mitochondrial localization and phenotypes linked to mitochondrial biology associated with loss of Mam33 (Muta et al., 1997; Feichtinger et al., 2017). We find here that Δ*mrx6* and Δ*mam33* share elevated mtDNA CN, but that Δ*mam33* shows a more severe growth defect than Δ*mrx6*. Moreover, more drastic changes on the mitochondrial proteome are observed in Δ*mam33* compared to Δ*mrx6* cells, pointing to Mrx6 independent roles of Mam33. Such roles could involve a chaperone function of Mam33 in ribosome assembly, which has recently been proposed (Hillman & Henry, 2019). Accordingly, the translation of Cox1 is significantly impaired in Δ*mam33* cells (Roloff & Henry, 2015). Therefore, Mam33 could be ideally positioned to coordinate mitochondrial translation, mtDNA replication and perhaps even transcription. Recruitment of Mam33 to unassembled ribosomal subunits would result in its detachment from Mrx6, which would be destabilized according to our data. Consequently, altered Pim1 and Cim1 activity and increasing Rpo41 levels could then impact mtDNA replication or transcription. However, it has to be noted that we did not observe greatly altered mitochondrial mRNA levels in Δ*mrx6* cells (Schrott & Osman, 2023).

Our current structural insights into a Pim1-Mrx6-Mam33 complex are primarily supported by AlphaFold predictions and mutational analyses that substantiate this interaction. Specifically, mutations of critical residues predicted to be involved in the Pim1-Mrx6 interaction destabilize the complex and result in increased mitochondrial DNA CN, thereby supporting our hypothesis that Mrx6 regulates mtDNA CN through its interaction with Pim1. Obtaining high-resolution structural information and *in vitro* reconstitution of the complex remain critical next steps. Such experiments will provide mechanistic insights into how Mrx6, in concert with Mam33, may modulate Pim1 activity.

Our investigation revealed an intriguing evolutionary perspective: Pim1 is highly conserved from yeast to human and also Mam33 has a human homolog, which when expressed in yeast rescues growth defects associated with Δ*mam33* strains (Muta et al., 1997). On the contrary, we have not identified homologs of the Pet20 domain family in higher eukaryotes. Our analysis uncovered that the Pet20 domain comprises a bipartite motif that comprises short characteristic stretches with only a few highly conserved residues. These characteristics combined with a potentially divergent nature of these motifs may explain the challenge in identifying homologous stretches in related proteins. The identification of a conserved glutamate residue (E479 in Pim1) that is present across diverse eukaryotes and participates in a critical interaction with the Pet20 motif, suggest potential functional conservation. Future research should focus on examining LonP protease interactors in higher eukaryotes to potentially identify functional homologs of the Pet20 domain containing proteins.

In summary, our analyses unveil a previously unrecognized organizational mechanism of the Lon protease, providing fundamental insights into substrate selection processes. Moreover, we have revealed a sophisticated regulatory network governing mitochondrial DNA CN, with the Lon protease functioning as a central mediator. These findings significantly advance our understanding of mitochondrial homeostasis and offer promising avenues for future investigations into the intricate molecular mechanisms regulating mtDNA CN.

## MATERIALS AND METHODS

### Yeast strains and growth conditions

Yeast strains were generated through stable integration of constructs into the haploid yeast genome based on homologous recombination and standard molecular biology techniques. Construction details and required plasmids are described in Supplementary Tables 1. Unless specified in addition to the listed strain numbers, clone A was used for final experiments when multiple confirmed clones were stocked. Strains were generated in the W303 (*leu2-3,112 can1-100 ura3-1 his3-11,15*) WT backgrounds yCO380 or yCO381 containing LacO repeats integrated in their mtDNA (Osman et al., 2015). Strains for Supplementary Figures S3H+L were generated in WT background yCO363, which carries an additional *ade2-1* mutation (Göke et al., 2020). Briefly, log phase cells were transformed by lithium acetate based transformation (transformation mixture for 4.5 OD cells washed with 0.1 M lithium acetate: 240 µl 50% (w/v) PEG3500, 34 µl H_2_O, 10 µl salmon sperm DNA (10 mg/ml), 36 µl lithium acetate (1M), 40 µl DNA) with linearized plasmids or PCR amplified fragments. Initial selection for deletions and integrations was performed by antibiotic resistance cassettes (Janke et al., 2004) or auxotrophic selection for an integrated *Kluyveromyces lactis URA3* gene flanked by hisG repeats allowing subsequent removal by 5-Fluoroorotic Acid Monohydrate (5-FOA) selection (Earley & Crouse, 1996). The generation of marker-less yeast mutants at the genomic locus of *MRX6* was based on a two-step selection process involving the counterselection of integrated *Kluyveromyces lactis URA3* with 5-FOA (Storici et al., 2001; Storici & Resnick, 2003). Successful strain generation was confirmed by colony PCR or sequencing of the respective locus. Most mutants were additionally backcrossed to minimize putative clonal genetic alterations or off-targets and reintroduce intact mtDNA. All depicted experiments were carried out with strains of mating type a, with the exception of mating type alpha strains in S3K,L,N-P. Cells were cultivated as published previously (Schrott & Osman, 2023). Liquid cultures were grown in yeast complete (YP) medium (1% yeast extract (VWR 84601), 2% bacto-peptone (VWR 26208), 0.004% adenine (all w/v), pH 5.5) containing either 2% (w/v) glucose (YPD) or 3% (v/v) glycerol (YPG) as carbon source with an agitation of 160 rpm at 30°C. For plates, medium was supplemented with 2% (w/v) bacto-agar and antibiotics were added for selection of transformants in the following concentrations: 300 µg/ml hygromycin, 300 µg/ml G418, 100 µg/ml nourseothricin. YPG plates containing additionally 0.1% glucose (indicator plates) were used to determine petite frequencies and perform tetrad dissections of P_Estr._-*Rpo41* strains. Synthetic defined (SD) selection plates without uracil of pH 5.5 contained 0.67% (w/v) yeast nitrogen base, 0.192% (w/v) dropout mix minus uracil (US Biological D9535), 2% (w/v) glucose and 2% (w/v) bacto-agar. For synthetic complete selection plates additionally 0.1% (w/v) 5-FOA and 0.005% (w/v) uracil were added. Sporulation plates contained 0.3% (w/v) potassium acetate, 0.048% (w/v) drop-out mix minus uracil, 0.00125% (w/v) uracil and 2% (w/v) bacto-agar.

### Drop dilutions growth analysis and determination of petite frequency

Growth analysis by drop dilutions and determination of petite frequencies was performed as described previously (Schrott & Osman, 2023).

### Isolation of genomic DNA and determination of mtDNA CN by efficiency-corrected quantitative PCR (qPCR)

Isolation of genomic DNA and determination of mtDNA CN by efficiency-corrected quantitative PCR (qPCR) was performed as described previously(Schrott & Osman, 2023). For Supplementary Figures S3L-N an earlier methodology without efficiency-correction using primers for *COX1* and *ACT1* was conducted (Göke et al., 2020).

### Denaturing total cell lysis

Denaturing total cell lysis for of at least 3 × 10^7^ mid-log phase grown cells was performed as described previously (Schrott & Osman, 2023). Cells were harvested for 4 min at 3146 g and washed dependent on amounts with ~0.5 to 1 ml of H_2_O. After 2 min centrifugation at 11,000 g pellets were stored at −80°C. Cells were quickly thawed and resuspended in 300 µl H_2_O per 7 × 10^7^ cells before the same amount of 0.2 M NaOH was added. The samples were incubated for 5 min at 25°C with repeated vortexing. Cells were collected at 3000 g and 4°C for 5 min and the pellet was resuspended in 150 µl of 1x Laemmli buffer (60 mM Tris/HCl pH 6.8, 2% w/v SDS, 8% v/v glycerol, 2.5% v/v *β*-mercaptoethanol, 0.0025% w/v bromophenol blue) per 7 ×10^7^ cells, frozen at −80°C and boiled for 5 min at 95°C. Undissolved components were pelleted for 1 min at 20,000 g, followed by the separation of 15 µl of the soluble sample (equal to 0.7 ×10^7^ cells) via Tris-glycine-SDS-PAGE.

### Immunoblot analysis

Immunoblot analysis was performed as published previously with minor modifications in blotting currents and durations (Schrott & Osman, 2023). Protein samples were separated by SDS-PAGE with self-casted 10 or 12% Tris-Glycine gels of 1 mm thickness and a width of 8.6 cm for 15 min at 80 V followed by ~1.5 h at 120 V. Proteins were transferred on PVDF membranes (Amersham Hybond P 0.2 PVDF, Cytiva 10600021) by semi-dry transfer in Towbin buffer (25 mM Tris, 192 mM Glycine, 0.05% (w/v) SDS, 20% (v/v) MeOH, pH 8.3 with NaOH). Blotting of 12% gels was performed for 60 min at a constant current of 1.5 mA/cm^2^ and membranes were subsequently cut and probed with antibodies against Pim1, Tim50, either Mrpl36 or Cim1, and Abf2. For probing of cut membranes with antibodies directed against Rpo41, Mrx6 and Mam33 10% gels were used and blotted for 90 min at 2 mA/cm^2^ to facilitate transfer and detection of the large ~150 kDa Rpo41. For panels 2G, 4H, 6F-K and S7G proteins were separated on 12% gels and blotted for 60 min at a constant current of 1.5 mA/cm^2^ for probing with antibodies directed against Mrx6, Mam33 and Rpo41. For 6B Towbin buffer lacked MeOH and all proteins were run on 10% gels and transferred for 90 min with 2 mA/cm^2^. The blotting duration was further increased to 150 min at 2 mA/cm^2^ for panels 5A-M and probing with antibodies against Rpo41, Mrx6, Cor2, Pim1 and Atp2 of S8C. For probing with antibodies against Cox2, Tim23, Mam33 and Cox17 of S8C, proteins were transferred for 90 min at 2 mA/cm^2^. The quality of semi-dry transfers was assessed by observation of the translucent protein bands that appeared during the drying process of the PVDF membranes. After completed transfer, the gels were subjected to colloidal Coomassie G-250 (C.B.B.) staining (Dyballa & Metzger, 2009) to control for uniform transfer, while the remaining high molecular weight protein bands retained in the gel after transfer served as additional loading control. The air-dried membranes were reactivated with MeOH, washed with TBS (50 mM Tris, 150 mM NaCl, pH 7.6), and blocked for at least 2 h in blocking buffer (TBS containing 5% (w/v) milk powder) at 25°C. Membranes were cut according to the sizes of the proteins and incubated overnight at 4°C rolling in 50 ml falcons with primary antibodies diluted in blocking buffer as listed in Supplementary Tables 1. Antibodies against Abf2, Cim1, Mam33, Mrx6, Pim1 and Rpo41 were used as previously described (Schrott & Osman, 2023). Several antibodies inherited from the Neupert lab were kindly provided by PD Dr. Dejana Mokranjac (Mrpl36 (Szyrach et al., 2003), Tim23 (C) (Popov-Čeleketić et al., 2008), Tim50 (IMS) (Mokranjac et al., 2003)), Dr. Nikola Wagener (Cor2, Cox2 (Cruciat et al., 1999), Cox17 (Mesecke et al., 2005)) and Dr. Max Harner (Atp2 (Harner et al., 2016)). After incubation with the primary antibody, membranes were washed in TBS for at least 30 min with three buffer exchanges at 25°C. A goat anti-rabbit (H + L)-HRP conjugated secondary antibody (Biorad 1706515) diluted 1:5000 in blocking buffer was added for at least 2 hours rolling at 25°C. Membranes were washed in TBS for at least 30 min with three buffer exchanges at 25°C, before chemiluminescent signals were detected with a Vilber Fusion FX imaging system, after a 1:1 mixture of HRP substrate solution 1 (100 mM Tris/HCl pH 8.5, 0.44 mg/ml Luminol (from 100x DMSO stock), 0.066 mg/ml p-coumaric acid (from 227 x DMSO stock)) and solution 2 (100 mM Tris/HCl pH 8.5, 0.18 µl/ml hydrogenperoxide) was applied to the blots. For detection of weak signals (Cim1, Rpo41 or Mrx6) SuperSignal™ West Atto Ultimate Sensitivity Substrate (Thermo Scientific A38554) was used. Chemiluminescence signals were quantified in Fiji/ImageJ (Schindelin et al., 2012) similar to previous reports (Stael et al., 2022). Measurements were performed on images obtained with identical exposure times for the data presented in panels 5A-M, 6B, S8C and S9A-C+F. For higher accuracy, blots of figure 6A were developed and analyzed with three different exposure times. First, normalization was performed for each exposure time relative to the intensity averaged from the three biological replicates of the controls cultivated with 10 nM estradiol. The normalized intensities were then averaged across all exposure times for the individual biological replicates (including each control), from which the overall mean and standard deviation were determined. A similar analysis was conducted for panels 6F-K, utilizing three exposure times for Cim1 and Rpo41, and two exposure times for Abf2, Pim1, and Tim50. Initially, the signals were calibrated for each exposure relative to a WT sample loaded, followed by normalization to Tim50.

### Isolation of mitochondria for immunoprecipitations

Mitochondria for immunoprecipitation experiments were isolated as decribed previously (Schrott & Osman, 2023) from 2 liters of log phase (undil. OD_660_ ~0.7) yeast cultures grown in YPG in 5-liter flasks, under ~150 rpm agitation at 30°C. Cells were harvested at 3993 g for 5 min and washed twice with 50 ml of H_2_O with a 2 min centrifugation at 3146 g. Cells were incubated for 10 min at 75 rpm agitation and 30°C in 2 ml/g wet weight of alkaline solution (100 mM Tris, 10 mM DTT) and pelleted as also in the following washing step at 1861 g for 5 min. After washing with 6.7 ml/g wet weight in spheroplast buffer (20 mM KH_2_PO_4_ pH 7.4, 1.2 M sorbitol), cells were gently resuspended in the same buffer containing 6/6.6 mg/ml Zymolyase (R) 20T (Amsbio 120491-1) to digest cell walls for 30 min at 30°C and 75 rpm agitation. Spheroplasts were then harvested for 5 min at 3146 g and 4°C and the buffer was exchanged without resuspension by pelleting cells again after addition of at least 10 ml of ice cold homogenization buffer (10 mM Tris/HCl pH 7.4, 0.6 M sorbitol, 1 mM EDTA, 0.2% BSA fatty acid free, 1 mM PMSF). Spheroplasts were broken with 15 cycles of dounce homogenization (Kimble^®^ 885300-0100, clearance 0.0005-0.0055 in.) in 6.8 ml/g wet weight homogenization buffer. Unbroken cells and nuclei were removed by 5 min centrifugation at 1861 g and 4°C. Mitochondria were collected from the supernatant for 10 min at 17,750 g and 4°C. Mitochondria were gently resuspended in SEM buffer (250 mM sucrose, 1 mM EDTA, 10 mM MOPS-KOH pH 7.2) containing 1x cOmplete™, EDTA-free protease inhibitor cocktail (Roche, Basel) and mitochondria were centrifuged for 10 min at 13,000 g and 4°C. The pellets were snap frozen in liquid nitrogen and stored at −80°C until use.

### Isolation and preparation of mitochondria for proteome analysis

A modified methodology was employed for the isolation of mitochondria for proteomic analysis. Yeast cultures cultivated to late log phase (undil. OD_660_ ~1) in 600 ml YPG in 2-liter flasks, under ~150 rpm agitation and at 30°C. Cells were harvested in a 500 ml centrifuge bottle in two centrifugation rounds at 2831 g for 10 min. Cells were washed once with 500 ml of H_2_O at 2831 g for 5 min. All cultures had a wet weight of ~2.3 g and identical steps according to the protocol above were performed until homogenization buffer was added. To the spheroplast pellet, 10 ml of ice cold homogenization buffer (10 mM Tris/HCl pH 7.4, 0.6 M sorbitol, 1 mM EDTA, 0.2% BSA fatty acid free, 1 mM PMSF) were added, initiating the flotation of the pellet via gentle agitation, without achieving complete suspension. An additional 10 ml of homogenization buffer was added and the spheroplasts were recollected for 5 min at 3146 g and 4°C. The supernatant was discarded, and the spheroplasts were gently opened in 15 ml of homogenization buffer through 15 vigorous pipetting cycles with a 1 cm cut-off 5 ml tip on a P5000. Samples were centrifuged for 5 min at 1861 g and 4°C. The supernatant was transferred to a separate tube and the homogenization and centrifugation steps were repeated with the remaining pellet. The combined 30 ml of supernatant was centrifuged for 5 min at 1861 g and 4°C. Mitochondria were collected from the supernatant for 10 min at 17,750 g and 4°C and gently resuspended in SEM buffer (250 mM sucrose, 1 mM EDTA, 10 mM MOPS-KOH pH 7.2) without protease inhibitor. Protein concentrations were determined by Bradford assay. 300 µl of mitochondria adjusted to a concentration of 1 µg/µl in SEM buffer were gently layered on top of 750 µl of SEM_500_ (500 mM sucrose, 1 mM EDTA, 10 mM MOPS-KOH pH 7.2) in 1.5 ml tubes, followed by cushion purification for 10 min at 13,000 g and 4°C. The supernatant and any loose, whitish portions of the pellet were removed from top to bottom. The resultant pellets were snap frozen in liquid nitrogen and stored at −80°C. Pellets were subjected to Trypsin digestion as reported previously (Wiśniewski et al., 2009).

### Spot-Immunoprecipitation from whole cell lysates

40 × 10^7^ mid-log phase yeast cells cultivated in 80 ml YPG in 300 ml flasks with an agitation of ~150 rpm at 30°C were harvested for 4 min at 3993 g. Cells were washed with 1 ml H_2_O and collected in 1.5 ml tubes for 2 min at 11,000 g. Pellets were flash frozen in liquid nitrogen and stored at −80°C until the next day. Screw cap 2 ml micro tubes (Sarstedt 72.694.406) were filled with a volume of an entire PCR tube (~250 µl) of 0.1 mm Zirconia beads (Roth N033.1) and precooled on ice. Cell pellets were quickly thawed on ice, resuspended by pipetting in 1 ml pulldown buffer (0.5% Nonidet P40, 15 mM Tris/HCl pH 7.4, 80 mM KCl, 2 mM EDTA pH 8.0, 0.2 mM spermine, 0.5 mM spermidine, 1x cOmplete™, EDTA-free protease inhibitor cocktail (Roche, Basel)) and transferred to the bead-equipped Screw cap tubes. Samples were ruptured 6 times for 30 s with 30 s break in between in a Speed-Mill PLUS 230 V (Analytik Jena 845-00007-2) homogenizer with −20°C precooled sample holder. Screw cap tubes were punched with a needle on the bottom to generate a hole and placed in 2 ml tubes. The liquid was separated from the Zirconia beads by centrifugation for 1 min at 1000 g and 4°C. Lysates were further cleared by centrifugation for 10 min at 20,000 g and 4 °C. 880 µl of the supernatant (input) were combined in 2 ml tubes with magnetically separated Spot-nanobody coupled M270 magnetic particles (ChromoTek Spot-Trap®Magnetic Particles M-270 etd-20) equivalent to 20 µl of bead slurry that has been pre-washed at least twice in a pooled manner for all samples with 1 ml of pulldown buffer. The samples were rotated over head for 1 h at 10 rpm and 4°C. Following the removal of the flow-through, magnetic particles were washed twice with 1 ml of pulldown buffer. Proteins were eluted with 46 µl of 1x Laemmli buffer (60 mM Tris/HCl pH 6.8, 2% w/v SDS, 8% v/v glycerol, 2.5% v/v *β*-mercaptoethanol, 0.0025% w/v bromophenol blue) for 5 min at 95°C. The samples were saparated via Tris-glycine-SDS-PAGE equivalent to 1% of input and flow-through, with 50% of the elution fractions loaded.

### Spot-Immunoprecipitation from mitochondrial lysates

Mitochondria were lysed on ice for a minimum of 15 min with two rigorous pipetting mixing steps in pulldown buffer (0.5% Nonidet P40, 15 mM Tris/HCl pH 7.4, 80 mM KCl, 2 mM EDTA pH 8.0, 0.2 mM spermine, 0.5 mM spermidine) supplemented with either 1x cOmplete™, EDTA-free protease inhibitor cocktail (Roche, Basel) (immunoblots 1C+E) or 2 mM of the serine protease inhibitor 4-(2-aminoethyl)-benzylsulphonyl fluoride hydrochloride (Pefabloc®) (immunoblots S2B+C and LC-MS/MS), to achieve a final concentration of 3 mg/ml (immunoblots 1C+E and S2B+C) or 3.5 mg/ml (LC-MS/MS 1D+F). The lysed samples were cleared from debris by centrifugation for 5 min at 3000 g and 4°C. Amounts equal to 2.5 mg (immunoblots 1C+E) or 3.5 mg (LC-MS/MS 1D+F) mitochondria were combined in 2 ml tubes with magnetically separated Spot-nanobody coupled M270 magnetic particles (ChromoTek Spot-Trap®Magnetic Particles M-270 etd-20) equivalent to 20 µl of bead slurry that had been pre-washed at least twice in a pooled manner for all samples with 1 ml of the respective pulldown buffer. For bead comparison in Supplementary Figures S2B+C, lysates were divided, and 417 µl (1.25 mg mitochondria) of lysate were in comparison also added to Spot-nanobody coupled magnetic agarose beads (ChromoTek Spot-Trap® Magnetic Agarose etma-20), both bead types corresponding to 10 µl of washed slurry per sample. The samples were rotated over head for 1 h 15 min (immunoblots) or 1 h 30 min (LC-MS/MS) at 10 rpm and 4°C. Following the removal of the flow-through, magnetic particles were washed twice with either 1 ml or 500 µl (S2B+C) of the pulldown buffer. Samples designated for LC-MS/MS analysis were further processed as described in the on-bead digest procedure. For immunoblots, proteins were eluted with 46 µl of 1x Laemmli buffer (60 mM Tris/HCl pH 6.8, 2% w/v SDS, 8% v/v glycerol, 2.5% v/v *β*-mercaptoethanol, 0.0025% w/v bromophenol blue) for 5 min at 95°C. The samples were saparated via Tris-glycine-SDS-PAGE equivalent to 0.6% (1C+E) or 1.2% (S2B+C) of input and flow-through, with 50% of the elution fractions loaded.

### HA-Immunoprecipitation from whole cell lysates

20 OD mid-log phase yeast cells cultivated in YPGal with an agitation of 160 rpm at 30°C were harvested for 5 min at 5000 g. Cell pellets were resuspended by pipetting in 500 µl lysis buffer (0.5% Triton X-100, 10 mM Tris/HCl pH 7.5, 150 mM NaCl, 0.5 mM EDTA pH 8.0, 1 mM phenylmethylsulfonyl fluoride (PMSF)) and transferred to screw cap tubes. Samples were ruptured using a FastPrep-24 5G homogenizer (MP Biomedicals, Heidelberg, Germany) with 3 cycles of 30 s, speed 8.0 m/s, 120 s breaks, glass beads) at 4°C. Lysates were cleared by centrifugation for 5 min at 16,000 g and 4°C. The supernatant was combined with 30 µl of Amintra Protein A Resin and 3 µl monoclonal Anti-HA antibody (Sigma 22190322). The bead slurry was pre-washed twice, pooled for all samples with 1 ml lysis buffer. The samples were rotated end-over-end for 1 h at 4°C. Following the removal of the flow-through, beads were washed twice with 500 µl of wash buffer II (50 mM Tris/HCl pH 7.5, 150 mM NaCl, 5% glycerol, 1 mM PMSF). Proteins were eluted with 50 l of 1x Laemmli buffer for 5 min at 95°C. The samples were separated via Tris-glycine-SDS-PAGE.

### On-bead digest of immunoprecipitations

After the two washing steps with pulldown buffer, detergent, protease inhibitors and salts were removed quickly at 4°C by two washes with 500 µl wash buffer 1 (15 mM Tris/HCl pH 7.4, 40 mM KCl, 2 mM EDTA), two washes with 500 µl wash buffer 2 (50 mM Tris/HCl pH 7.4) and one wash with 100 µl wash buffer 3 (50 mM Tris/HCl pH 8.0). To each sample, 80 µl of freshly prepared Trypsin buffer (50 mM Tris/HCl pH 8.0, 2 M Urea, 1 mM DTT, 0.3/80 µg/µl Trypsin/Lys-C ((Promega V5071) from freshly prepared 1 µg/µl stock in 50 mM acetic acid)) containing 0.3 µg Trypsin/Lys-C were added and proteins were digested for 100 min (Pim1-Spot in WT and Δ*mrx6*) or 110 min (Mrx6-Spot and WT) at 37°C. For all incubation steps samples were agitated a 500 rpm and a heating lid was used to prevent condensation. Supernatants were transferred to a fresh tube and beads were washed twice with 60 µl of wash buffer 4 (50 mM Tris/HCl pH 8.0, 1 M Urea) collecting all supernatants in one tube. Samples were reduced with 0.72 µl of 1 M DTT (4 mM final concentration: 3.6 mM extra to remaining 0.4 mM from Trypsin buffer) for 30 min at 37°C. For subsequent alkylation 10.5 µl of freshly dissolved 200 mM Iodoacetamide (Promega VB1010) were added (final concentration of 10 mM) and samples were incubated for 30 min at 25°C. Additional 0.2 µg of Trypsin/Lys-C (2 µl of freshly 1:10 diluted 1 µg/µl stock solution in 50 mM acetic acid to 0.1 µg/µl in 50 mM Tris/HCl) were added for overnight digest at 25°C (13.5 h for (Pim1-Spot in WT and Δ*mrx6* and 12.5 h for Mrx6-Spot and WT). The digest was stopped by addition of 5.62 µl of 50% formic acid (1.33% final concentration) and 3 µl of the sample were controlled on pH indicator paper (pH 2-3). Samples were centrifuged for 1 min at 4°C and 20000 g and shortly frozen at −20°C before peptides were desalted with home-made C18 stage tips (Rappsilber et al., 2003), vacuum dried and stored at −80°C.

### LC-MS/MS analysis

Liquid chromatography-tandem mass spectrometry (LC-MS/MS) was performed as described previously (Marino et al., 2019) on a nano-LC system (Ultimate 3000 RSLC, ThermoFisher Scientific, Waltham, MA, USA) coupled to an Impact II high-resolution Q-TOF (Bruker Daltonics, Bremen, Germany) using a CaptiveSpray nano electrospray ionization (ESI) source (Bruker Daltonics). The nano-LC system was equipped with an Acclaim Pepmap nano-trap column (C18, 100 Å, 100 µm × 2 cm) and an Acclaim Pepmap RSLC analytical column (C18, 100 Å, 75 µm × 50 cm), both from ThermoFisher Scientific. The peptide mixture was separated for 90 min over a linear gradient of 4-45% (v/v) Acetonitrile at a constant flow rate of 250 nl/min. The column was kept at 50°C throughout the run. MS1 spectra with a mass range from m/z 200-2000 were acquired at 3 Hz, and the 18 most intense peaks were selected for MS/MS analysis using an intensity-dependent spectrum acquisition rate of between 4 and 16 Hz. Dynamic exclusion duration was 0.5 min. Raw files were processed using the MaxQuant software v2.4.9.0 for immunoprecipitation samples and v1.6.10.43 for mitochondria samples (Cox & Mann, 2008). Peak lists were searched against the *Saccharomyces cerevisiae* (strain ATCC 204508/S288c) reference proteome (Uniprot, www.uniprot.org). For immunoprecipitations Mrx6-Spot, Pim1-Spot and the BC2 nanobody were included (Braun et al., 2016). A false discovery rate of 1% was applied at peptide and protein level. Proteins were quantified using the label-free quantification (LFQ) algorithm(Cox et al., 2014) with default settings. Enzyme specificity was set to Trypsin, allowing up to two missed cleavages. Cysteine carbamidomethylation was set as a fixed modification. Acetylation of protein N-termini and methionine oxidation were set as variable modifications. Statistical analysis was performed using Perseus (Tyanova et al., 2016) and R (R Core Team, 2019). Potential contaminants, reverse hits, and proteins identified only by site modification were excluded. Replicates were grouped and LFQ intensities were stringently filtered to contain valid values in all three replicates in at least one group for immunoprecipitation samples and in at least three of four replicates in at least one genotype for mitochondria samples. Protein LFQ intensities were log_2_-transformed and missing values were imputed from a normal distribution distribution with a width of 0.3 and a down-shift of 1.8 standard deviations to enable statistical evaluation by a two-tailed homoscedastic t-test with p-values corrected by the Benjamini-Hochberg approach (Benjamini & Hochberg, 1995) at a false discovery rate of 0.05. For immunoprecipitations the few comparisons affected by imputation were examined to exclude potential artifacts, as detailed in Supplementary Tables 2. Especially imputed values diminished actual fold enrichments for the most robust differences in immunoprecipitiations, when high intensities of proteins were exclusively detected in all replicates of one group, while remaining undetected in the comparative group, as confirmed by immunoblots. Cytosolic ribosomal contaminants were removed from the datasets. Statistical analysis for mitochondria proteomics was done in R using the Limma package (Ritchie et al., 2015). P-values were adjusted for multiple comparisons according to the Benjamini-Hochberg approach (FDR = 0.05) (Benjamini & Hochberg, 1995). For analysis of immunoprecipiations, on-bead digested peptides from three technical replicates of different mitochondrial aliquots derived from the same mitochondria isolation of each strain were analyzed. For analysis of the mitochondrial proteome, two biological independent mitochondrial isolates (for Δ*mrx6* two independently backcrossed strains) in two technical replicates each were used.

### Determination of nuclear transcript levels

Nuclear transcript levels were determined from total RNA isolated from 40 × 10^7^ cells cultured to log phase at 30°C in either YPD or YPG medium. RNA isolation, reverse transcription, and qPCR analysis was performed as described previously (Schrott & Osman, 2023).

### Estradiol regulated gene expression

Similar as reported for the modulation of *CIM1* or *PIM1* expression (Schrott & Osman, 2023), strains were generated to modulate the expression of *RPO41* or *MRX6* directly at the endogenous gene locus by different estradiol concentrations. The plasmid pSS037 facilitated the stable integration at the yeast LEU2 locus of an estradiol responsive transcriptional activation element, composed of a fusion of the LexA protein with an estrogen receptor ligand binding domain and the B42 transcriptional activator, regulated by an *ACT1* promoter and *CYC1* terminator (Ottoz et al., 2014). This construct was also expressed in control cells. The pSS037 encoded auxotrophic *Kluyveromyces lactis URA3* selection marker flanked by hisG repeats (Earley & Crouse, 1996), was removed from all strains employed in final experiments through selection with 5-Fluoroorotic Acid Monohydrate (5-FOA). The 5’ UTRs of *RPO41* or *MRX6* were modified by standard genomic integration using the Janke S1 as well as S4 primer binding sites (Janke et al., 2004), which flank the LexA-responsive promoter elements encoded on pSS045 (*kanMX4* selection cassette for P_Estr._-*RPO41*) or pSS046 (*natNT2* selection cassette for P_Estr._-*RPO41*). Similar to P_Estr._-*PIM1* strains, P_Estr._-*RPO41* strains were generated in the presence of 30 nM estradiol, backcrossed against WT *RPO41* backgrounds to refresh mtDNA, predominantly kept as cryo stocks and inoculated from freshly streaked YPG plates containing 10 nM estradiol. For all experiments involving P_Estr._-*PIM1* and P_Estr._-*RPO41*, log phase liquid YPG cultures with 10 nM estradiol (mimicking WT expression levels) were used as starting material before estradiol concentrations were modified. In all experiments, *β*-estradiol (Alfa Aesar L03801), stored as 1 mM ethanol stock at −20°C, was freshly diluted in medium to reach low final nM concentrations.

### Cycloheximide chase

For protein stability chase experiments following the inhibition of cytosolic translation by cycloheximide, WT and Δ*mrx6* cells were cultured overnight in YPG at 30°C until reaching stationary phase (OD_600_ ~2-4). Cultures were diluted to an OD_600_ of 0.2 in either 100 ml (4-hour chases) or 60 ml (1-hour chases) of YPG and grown to mid-log phase (OD_600_ ~0.4-0.6) for approximately 4 hours. For the comparison of different Pim1 levels, P_Estr._-*PIM1* cells (Schrott & Osman, 2023) were cultured to log phase throughout the day in YPG with 10 nM estradiol. Cells were washed with YPG, inoculated to 0.01 × 10^7^ cells/ml in 60 ml of YPG containing either 10 nM estradiol (WT-like Pim1 protein levels) or without estradiol (Pim1 downregulation), and cultured to mid-log phase for ~18 hours. Cycloheximide was added to all cultures at a final concentration of 100 µg/ml from freshly prepared ethanol stocks of 10 mg/ml (regular chases) or 20 mg/ml (Pim1 estradiol regulation). Equivalent cell volumes were rapidly harvested at 3146 g for 4 min at 1-hour (4-hour chases) or 15-minute intervals (1-hour chases) corresponding to ~7 × 10^7^ cells (4-hour chases and 1-hour chases with Pim1 regulation) or ~5 × 10^7^ cells (regular 1-hour chases). Cells were washed with 1 ml of H_2_O and immediately stored at −80°C. Subsequent denaturing total cell lysis was performed with adjusted concentrations in Laemmli buffer according to optical densities measured at at each chase start: 0.05 × 10^7^ cells/µl (regular 4- and 1-hour chases) or 0.07 × 10^7^ cells/µl (1-hour chase with Pim1 regulation). A sample volume of 10 µl was separated via Tris-glycine-SDS-PAGE and analyzed by immunoblotting. In graphs, the quantified immunoblot signals are presented as means (dots with black outline) ± SD, and separate biological replicates are represented as circle (#1), square (#2), triangle (#3) or diamond (#4) without outlines.

### Protein structure prediction and analysis

The protein sequences utilized for structure prediction refer to the *Saccharomyces cerevisiae* W303 reference genome and were obtained from the Saccharomyces Genome Database (http://www.yeastgenome.org). AlphaFold v2.2.4 (Jumper et al., 2021) was used for all structure predictions apart from the large assembly of 6x Pim1 NTD + ATPase, 3x Mrx6 and 3x Mam33. The following versions of model parameters and genetic databases were used: AlphaFold params, PDB_seqres, PDB_mmcif, MG-nify (as of December 19, 2024), UniRef90 (as of December 18, 2024), Uniclust30, UniProt (as of December 15, 2024), BFD (as of October 25, 2022) and PDB70 (as of October 12, 2022). Three seeds were run per model and relaxation was performed on all of the resulting 15 structure predictions. Models were ranked and selected based on confidence metrices: for the “monomer_ptm” mode prediction of Mrx6 the mean pLDDT confidence, and for the “multimer” mode predictions the ipTM + pTM scores. AlphaFold 3 (Abramson et al., 2024) was used via the AlphaFold Server (https://alphafoldserver.com) on May 17, 2024 to predict the structure of 6x Pim1 NTD + ATPase (181-770 Δ285-365), 3x Mrx6 (107-524) and 3x Mam33 (47-266), selecting the best-ranked model also according to the ipTM + pTM confidence. All structures were visualized utilizing ChimeraX 1.7 (Goddard et al., 2018; Meng et al., 2023). Additional build-in AlphaFold commands facilitated the visualize error estimates. Residue specific pLDDT confidences were color coded on the ribbon structures. Predicted Aligned Error (PAE) values were visualized through two different approaches: color-based clustering within multimer ribbon structures, and heat map plots for each pair of residues. Hydrogen bond pairs were also extracted using ChimeraX 1.7, with a distance tolerance of 0.4 Å and an angle tolerance of 20°. In the visualizations, hydrogen bonds meeting strict criteria of a distance at 3.5 ≤ Å and an angle near 180° were denoted by blue dashes, whereas those with relaxed criteria were represented in orange. The extracted hydrogen bonds between the Pet20 domain proteins and the Pim1 NTD were filtered for those that consistently appeared in the three highest-ranked predictions, representing either the N_Pet20_- or C_Pet20_-motif interactions, except for the N_Pet20_-motif interactions of Pet20, where in total only two models were predicted, in contrast to 13 for the C_Pet20_-motif. For hydrogen bonds between Mrx6 and Pim1 in the AlphaFold 3 structure of 6x Pim1 NTD + ATPase, 3x Mrx6 and 3x Mam33, the pairs also had to be present at all three reciprocal interaction sites of either obtuse- or acute angeled protomers.

### Phylogenetic analysis

Probabilistic consensus logos were constructed by inputting full-length homologous protein sequences into Skylign (Wheeler et al., 2014). The tool was set to create a Hidden Markov Model (HMM) by running HMMER3 version 3.1b1 (February 2013) (Eddy, 2011), remove mostly empty columns, handle alignment sequences as full length and estimate the letter height based on all information content. The residues’ colors were manually adjusted in accordance with their physico-chemical properties, utilizing the Jalview (Waterhouse et al., 2009) Zappo color scheme. For the Pet20 domain family, a comparison was made of 232 proteins classified under IPR014804/PF08692 from the InterPro/PFAM databases, while excluding the highly divergent sequences A0A3M7MRL5 and A0A1E4T1F1 (as of January, 2024). For the Pim1 consensus logo and alignments, protein sequences from all 34 reviewed eukaryotic chloroplastic/mitochondrial Lon protease homologues, categorized under InterPro IPR027503 (as of January, 2024), were utilized. Progressive-iterative alignments with full length protein sequences were conducted utilizing MUSCLE (Edgar, 2004) through the EMBL-EBI service tool (Madeira et al., 2022), then filtered and visualized with Jalview (Waterhouse et al., 2009).

### Data representation, statistics and visualization

In the manuscript, bar and line graphs illustrate biological replicates as discrete data points, while means are indicated as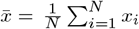, with the calculation of the sample standard deviation as 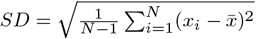 provided as bidirectional error bars. Two-tailed heteroscedastic t-tests were performed to assess the differences among biological replicates across datasets involving efficiency-corrected qPCR, petite frequency analysis, and quantified immunoblot chemiluminescence signals. The resulting p-values are represented as follows: n.s. p *>* 0.05, * p ≤ 0.05, ** p *<* 0.01, *** p *<* 0.001, **** p *<* 0.0001. The analysis, calculations, and generation of graphs for numerical data were conducted utilizing Microsoft Excel and Python 3. Sequence logos were generated using Skylign (Wheeler et al., 2014), and alignments were visualized with Jalview (Waterhouse et al., 2009). Immunoblot images were cropped to achieve uniform sizes and precise alignment to marker bands using IrfanView. For enhanced visualization of some images, the entire histograms were slightly adjusted by modifying the upper and lower limits of depicted pixel intensities in Fiji/ImageJ (Schindelin et al., 2012). Protein structures were visualized using ChimeraX 1.7 (Goddard et al., 2018; Meng et al., 2023). Figures were assembled and labeled using Affinity Designer v1.10.6.

## Supplementary

Supplementary Data of the manuscript are attached in separate files.

## Acknowledgements

We are grateful to the following colleagues, who kindly provided various antibodies: Prof. Dr. Dejana Mokranjac (Mrpl36, Tim23 (C) and Tim50 (IMS)), Dr. Nikola Wagener (Cor2, Cox17 and Cox2) and Dr. Max Harner (Atp2). We thank Elisa Adani for contributing as a Master’s student in early steps of the project, which influenced the final directions of the study. We extend our gratitude to Nadja Lebedeva and Tanja Kautzleben for assisting in preparation of plates and media and general lab management. We thank Sylvia Berngehrer and Tatiana Schubert for dish washing and autoclave service. In addition, we thank Nima Esmaeelpour for installing and updating AlphaFold v2.2.4. We further thank the ‘Mito-Club’ for stimulating discussions.

## Author contributions

Sim.S. designed, performed and analyzed experiments and prepared figures. L.B. performed pull-down experiments of catalytically inactive Pim1. C.G., S.B and R.R. assisted in experimental work under the supervision of Sim.S.. G.M. and Ser.S. carried out mass spectrometry analysis. Sim.S. and C.O. conceptualized the study. C.O. and J.M.H. supervised the study and provided funding. Sim.S. and C.O. wrote and edited the manuscript, with all authors critically reviewing it.

## Data availability

The datasets generated and analyzed during the current study are available from the corresponding author upon request. These include additional primer sequences employed in strain generation and verification, plasmid maps, imaged agarose gels, qPCR data, yeast plate images, petite frequency quantifications, immunoblots, predicted protein structure models, LC-MS/MS data, and further graphical analyses stored in Excel files and Python scripts.

