## Supplementary Figures for "Mrx6 binds the Lon protease Pim1 N-terminal domain to confer selective substrate specificity and regulate mtDNA copy number"

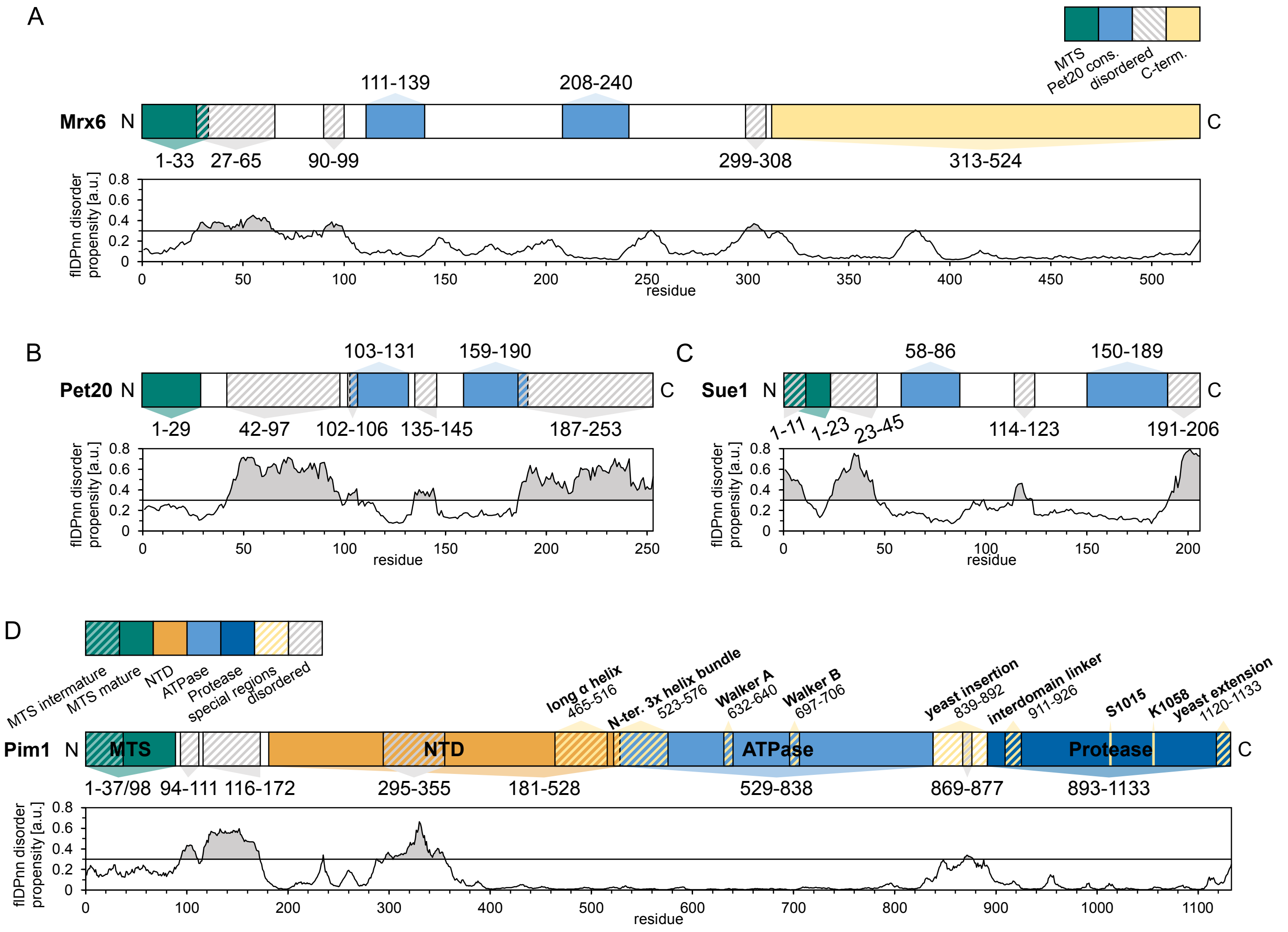


**Figure S1. Schematics of protein domains of Mrx6, Pet20, Sue1 and Pim1.**
Related to Figures 2 and 3.
(**A-D**) Characteristics of Mrx6 (A), Pet20 (B), Sue1 (C) and Pim1 (D) including depiction of the predicted flDPnn disorder propensity for each position (Hu et al., 2021). Disordered regions above the flDPnn threshold are highlighted with dashed lines. The MTS (mitochondria targeting sequence) is colored in dark green, the $N_{Pet20}$- and $C_{Pet20}$-motif of Mrx6, Pet20 and Sue1 in blue and the specific C-terminal extension of Mrx6 in ivory. (D) In addition to the Pim1 NTD (N-terminal domain) (orange), ATPase (blue) and protease (dark blue) domains, further special regions are labeled and highlighted with dashed yellow lines.

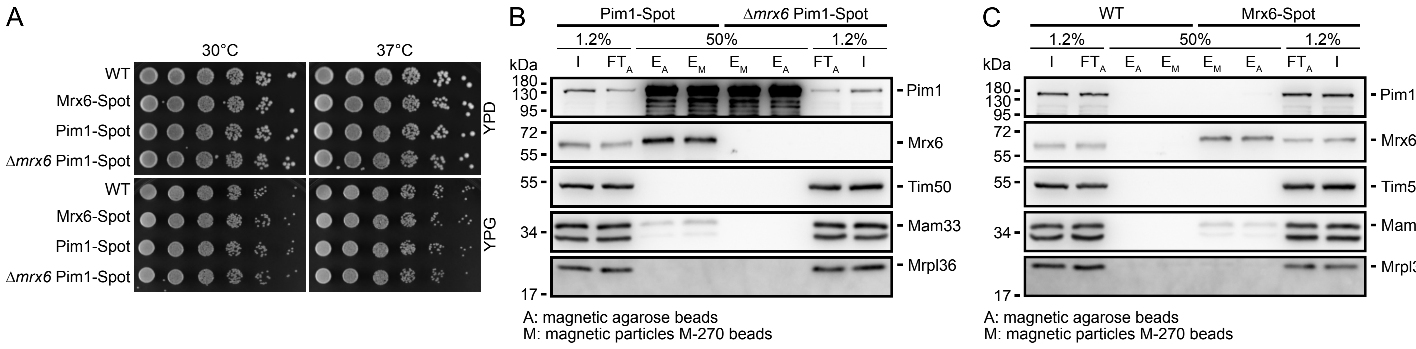


**Figure S2. Mrx6-dependent association between Mam33, Pet20 and Pim1.**
Related to Figure 1.
(**A**) Drop dilution growth analysis at 30 or 37°C of indicated strains pre-grown in YPD at 30°C. Images were taken after 48 h growth on YPD or YPG (**B+C**) Immunoblots of Pim1, Mrx6, Tim50 (control), Mam33 and Mrpl36 derived from M-270 ($_M$) or magnetic agarose ($_A$) coupled Spot nanobody immunoprecipitations of lysates from 1.25 mg isolated mitochondria of Pim1-Spot and Δ*mrx6* Pim1-Spot (B) or WT and Mrx6-Spot (C) cells cultivated in YPG at 30°C. Pim1 activity was inhibited by 2 mM Pefabloc ®. Input (I) and flow-through (FT) fractions represent 1.2% of total amounts and bound fractions eluted with 1x laemmli buffer ($E_{A/M}$) are 50% of the eluted volumes.

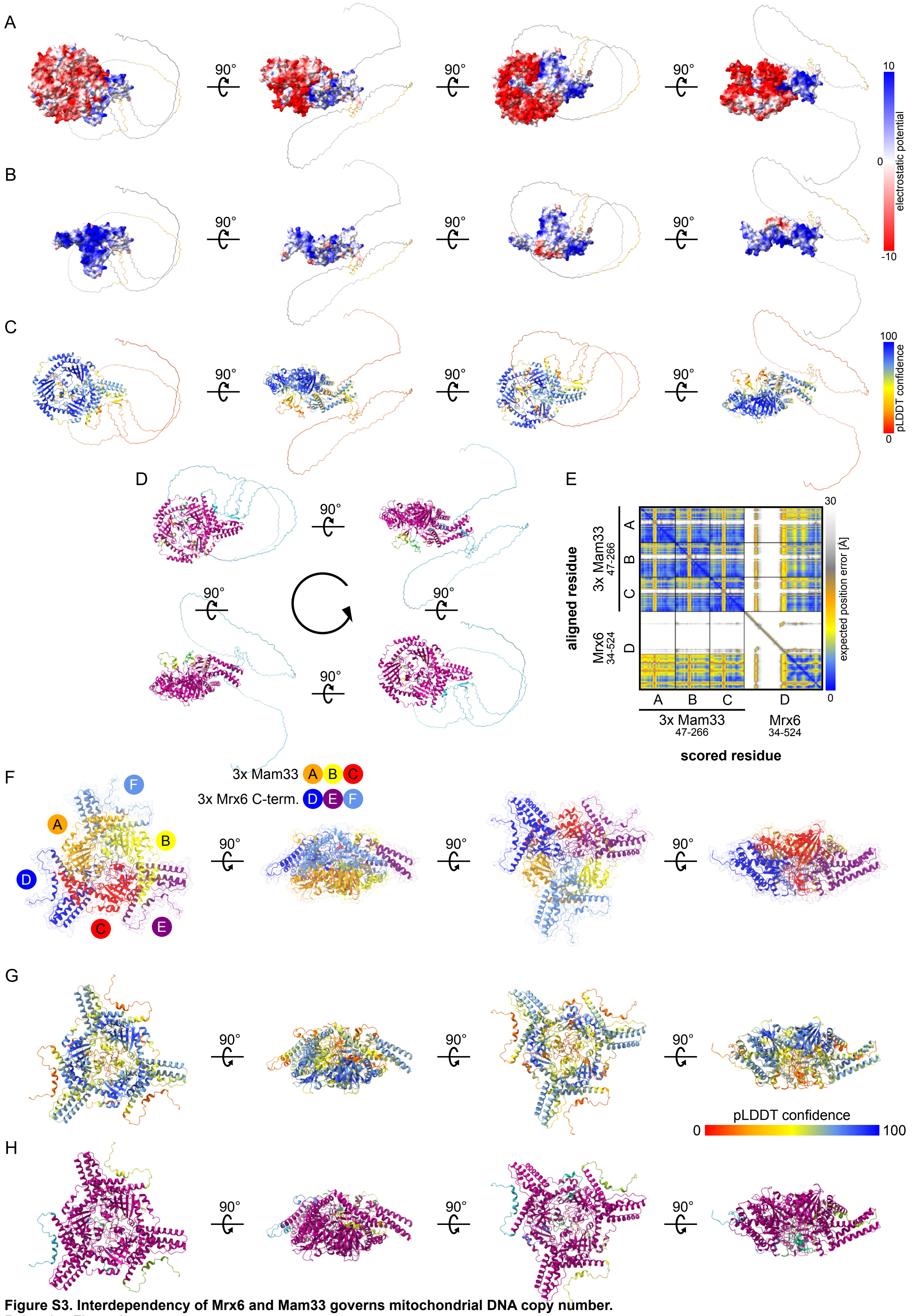


**Figure S3. Interdependency of Mrx6 and Mam33 governs mitochondrial DNA copy number.**
Related to Figure 2.
see full legend on next page.

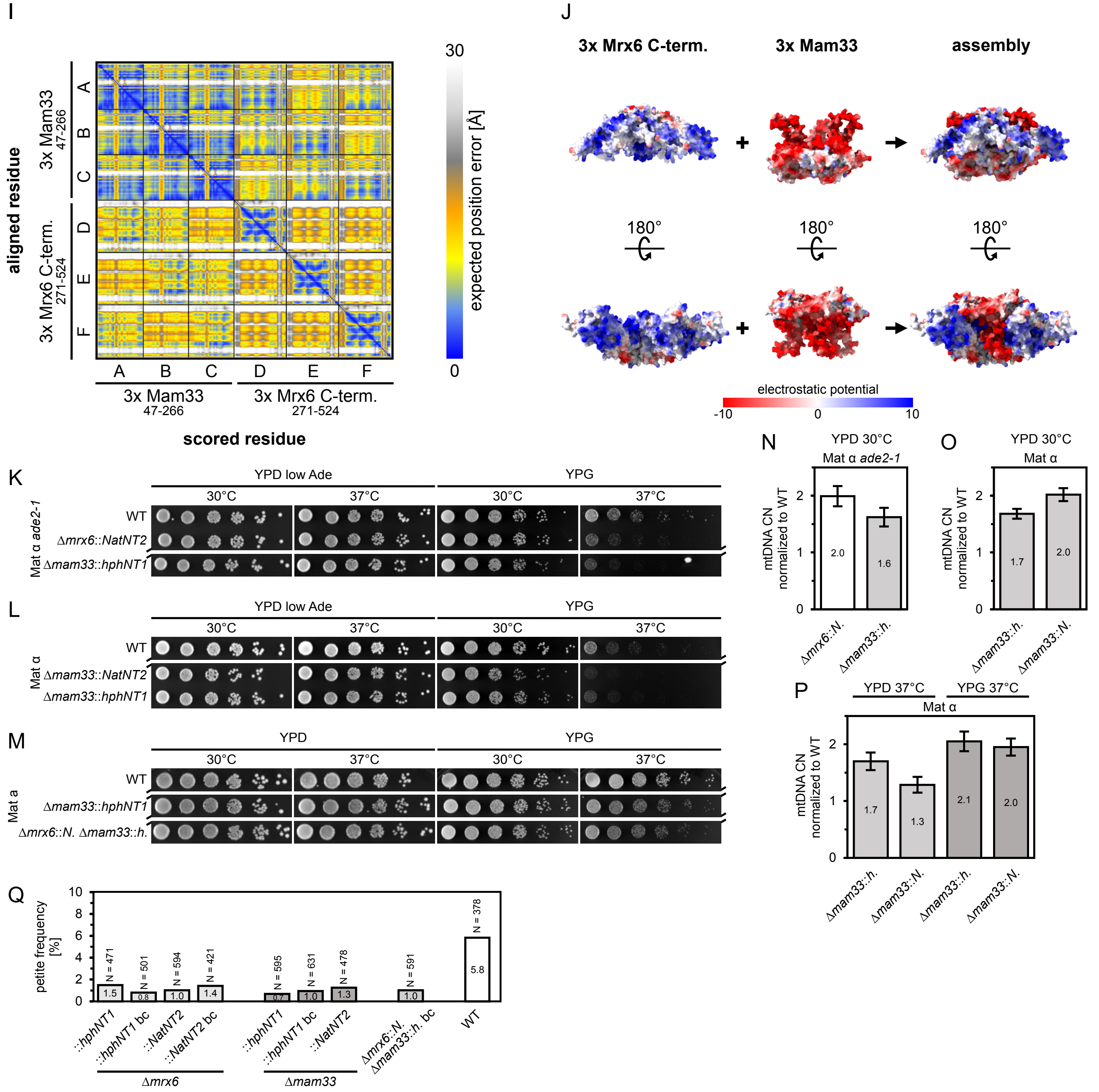


**Figure S3 Interdependency of Mrx6 and Mam33 governs mitochondrial DNA copy number.**
Related to Figure 2.
(**A-E**) AlphaFold multimer prediction of 3x Mam33 (residues 47-266) with one molecule of full length Mrx6 (residues 34-524). Best ranked model with ipTM + pTM confidence of ~0.8349 is depicted. The Mrx6 N-terminus is depicted as gray ribbon with Pet20 motifs highlighted in orange. The surface structures colored according to the coulombic electrostatic potential in the range of -10 to 10 are shown simultaneously for Mam33 and the Mrx6 C-terminus (A) or without Mam33 (B). Error estimates are shown on the ribbon structure colored according to the residue specific pLDDT confidence (C), separate coloring of coherent domains based on residue clustering according to the predicted aligned (PAE) value (D) or are depicted as a heat map plot of the expected position error for every pair of residues (E). (**F-J**) AlphaFold multimer prediction of 3x Mam33 (residues 47-266) with 3x Mrx6 C-terminus (residues 271-524). Best ranked model with ipTM + pTM confidence of ~0.7674 is depicted. Individual chains colored differently are shown as ribbons with transparent surfaces (F). Error estimates are shown on the ribbon structure colored according to the residue specific pLDDT confidence (G), separate coloring of coherent domains based on residue clustering according to the predicted aligned (PAE) value (H) or are depicted as a heat map plot of the expected position error for every pair of residues (I). In addition to top and bottom views in figure 2E, (J) shows side views of surface structures colored according to the coulombic electrostatic potential in the range of -10 to 10. (**K-M**) Drop dilution growth analysis at 30°C or 37°C of indicated strains pre-grown in YPD. Images were taken after 48 h growth on YPD (in H+I YPD with lower adenine concentration of 12 mg/l corresponding to 30 % of usual amount) or YPG. Two small black diagonals between cropped pictures indicate that these samples originated from the same experiment and plate and unrelated mutants of the experiment in between were removed for depiction. (**N-P**) qPCR on mtDNA encoded *COX1* and nuclear reference *ACT1* of different deletion strains of *MRX6* and *MAM33* grown under the indicated conditions. Data was normalized on the respective WT of the same genetic background from the same experiment. n = 1, data represent technically derived mean ± SD including error propagation of pipetting replicates of DNA samples. (**Q**) Petite frequencies of various Δ*mrx6* and Δ*mam33* strains as well as a Δ*mrx6*Δ*mam33* double mutant grown in YPD at 30°C. bc indicates that strains were backcrossed. n=1.

A

| combination | highest confidence | Lon interaction $N_{Pet20}$-motif | Lon interaction $C_{Pet20}$-motif | no interaction | total |
|---|---|---|---|---|---|
| Pim1 NTD (181-528) + Mrx6 N-term. (107-280) | 0.6575310 | 1 | 2 | / | 3 |
| Pim1 NTD (181-528) + Mrx6 (107-524) | 0.5886741 | / | 1 | 2 | 3 |
| Pim1 NTD (181-528) + Mrx6 (107-524) and Pet20 (30-253) | 0.4098974 | 1 (only Mrx6) | 2 (only Mrx6) | / | 3 |
| 2x Pim1 NTD (181-528) + Mrx6 (107-524) and Mam33 (64-266) | 0.2864556 | / | / | 15 | 15 |
| Pim1 NTD+ATPase (181-910) + Mrx6 (107-524) | 0.3484105 | 1 | / | 2 | 3 |
| Pim1 NTD+ATPase (181-910) + Pet20 (30-253) | 0.3881263 | 3 | / | / | 3 |
| Pim1 NTD (99-528) + Mrx6 N-term. (34-280) | 0.5315225 | 1 | 2 | / | 3 |
| Pim1 NTD (168-528) + Mrx6 (107-524) and 3 x Mam33 (47-266) | 0.5944092 | / | / | 15 | 15 |
| Pim1 NTD (99-528) + Pet20 (30-253) | 0.6080002 | / | 3 | / | 3 |
| Pim1 NTD (168-528) + Mrx6 N-term. (34-280) | 0.6648306 | 6 | 9 | / | 15 |
| Pim1 NTD (168-528) + Pet20 (30-253) | 0.6750746 | 2 | 13 | / | 15 |
| Pim1 NTD (168-528) + Sue1 (25-206) | 0.6972060 | 8 | 7 | / | 15 |
| LONP1 NTD (103-408) + Mrx6 N-term. (34-280) | 0.6823463 | / | 4 | 11 | 15 |
| LONP1 NTD (103-408) + Pet20 (30-253) | 0.4443096 | 1 (13 close) | / (1 close) | 14 | 15 |
| LONP1 NTD (103-408) + Sue1 (25-206) | 0.4519251 | / | / | 15 | 15 |

0.4 model confidence rank 0 1
ipTM + pTM

B

| combination | Lon interaction with $N_{Pet20}$-motif | | | Lon interaction with $C_{Pet20}$-motif | | |
|---|---|---|---|---|---|---|
| prediction | #1 | #2 | #3 | #1 | #2 | #3 |
| Pim1 NTD (168-528) + Mrx6 N-term. (34-280) | 0.6582350 | 0.6404202 | 0.6250581 | 0.6648306 | 0.6561704 | 0.6456119 |
| Pim1 NTD (168-528) + Pet20 (30-253) | 0.6540378 | 0.6475854 | / | 0.6750746 | 0.6724893 | 0.6698457 |
| Pim1 NTD (168-528) + Sue1 (25-206) | 0.6972060 | 0.6927281 | 0.6878195 | 0.6964365 | 0.6958779 | 0.6902738 |
| LONP1 NTD (103-408) + Mrx6 N-term.(34-280) | / | / | / | 0.6823463 | 0.6462255 | 0.5986310 |

0.4 model confidence rank 0 1
ipTM + pTM

**Figure S4. The Pet20 domain family proteins Mrx6, Pet20 and Sue1 are predicted to interact with the Pim1 NTD either via the $N_{Pet20}$-motif or the $C_{Pet20}$-motif, while the predicted human LONP1 interaction with Mrx6 is restricted to the $C_{Pet20}$-motif and absent for Pet20 or Sue1.** Related to Figure 3.
(**A**) Table of performed AlphaFold multimer predictions involving Mrx6, Pet20, Sue1, Mam33 and the Lon proteases Pim1 or LONP1. Either the first three highest ranked or all 15 predicted models were analyzed for the listed interactions between Pet20 motifs and the Lon proteases by coloring coherent domains in the multimer structures based on residue clustering according to the predicted aligned (PAE) value in ChimeraX 1.7 (Meng et al., 2023). (**B**) Table on ipTM + pTM confidences of indicated AlphaFold multimer predictions of Pet20 domain proteins and the Lon proteases Pim1 or LONP1. Values of the three highest ranked models in conformations where either the $N_{Pet20}$-motifs or the $C_{Pet20}$-motifs interact with the Lon protease are shown. These models were used for the prediction of hydrogen bond pairs summarized in Figure 3B and Supplementary Figures S5A-C.

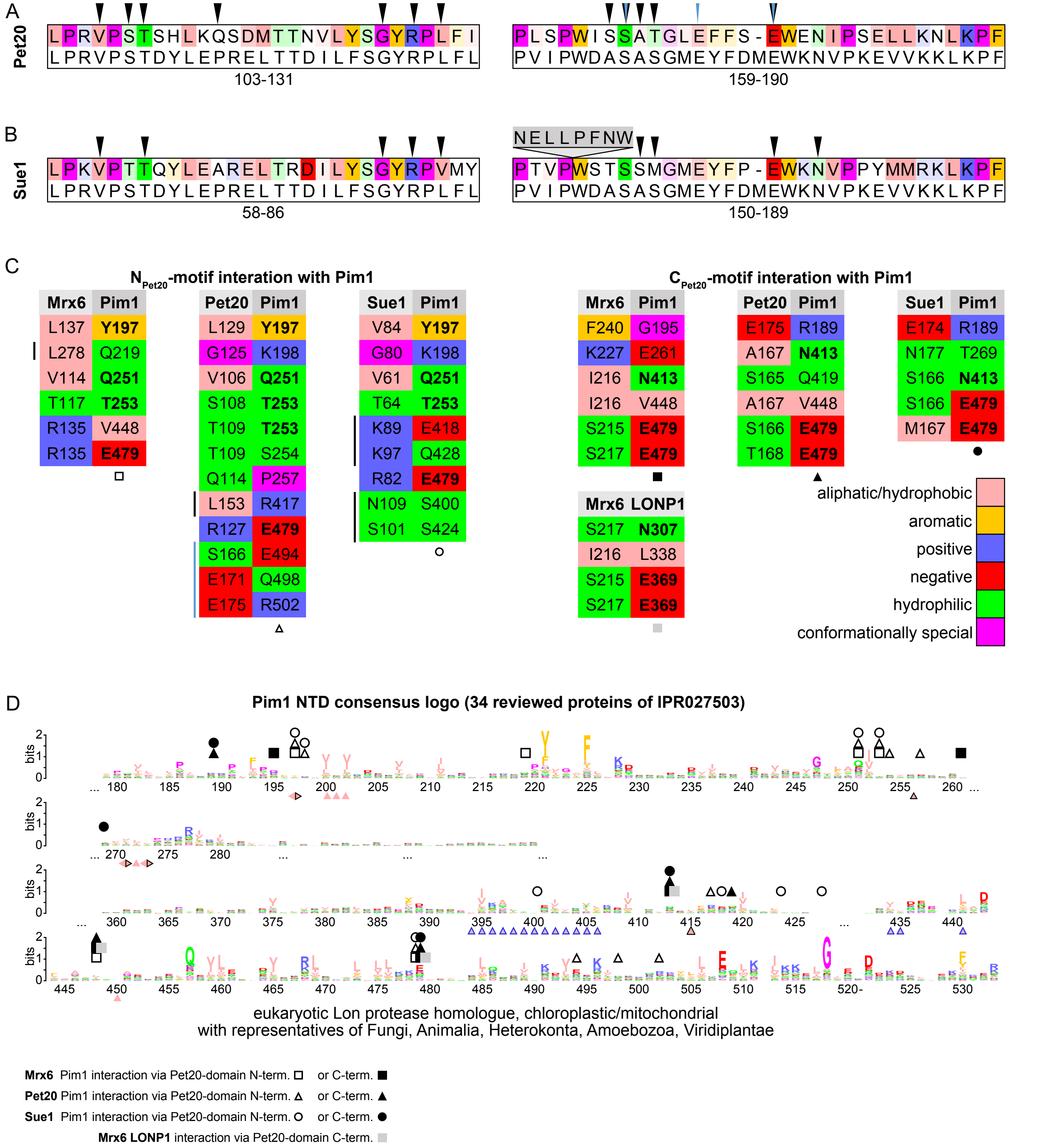


**Figure S5. Conserved motifs present in the Pet20 domain protein family interact with a conserved region of the Pim1 N-terminal domain.** Related to Figure 3.
see full legend on next page.

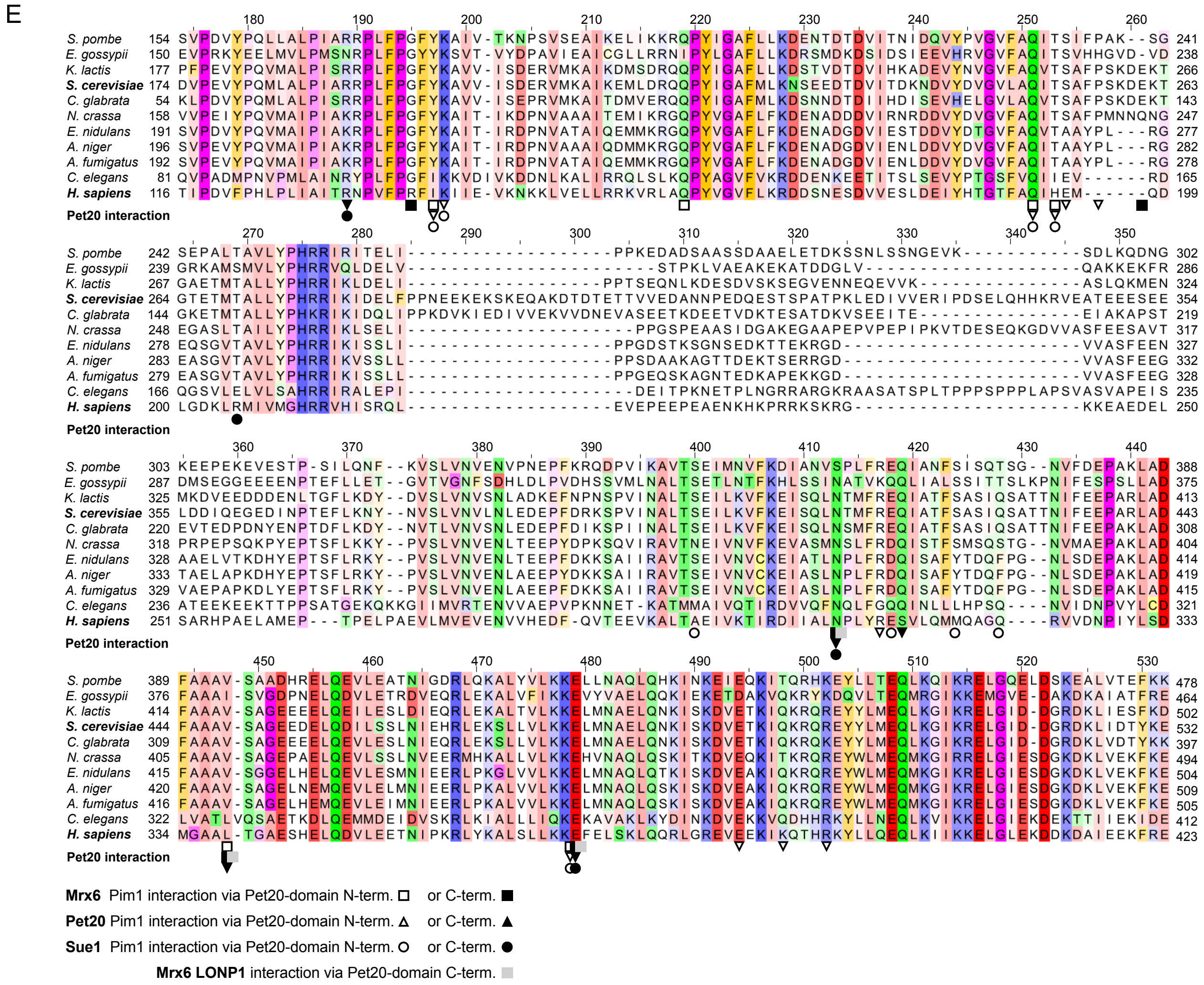


**Figure S5. Conserved motifs in the Pet20 domain protein family interact with a conserved region of the Pim1 N-terminal domain.**
Related to Figure 3.

(**A+B**) $N_{Pet20}$- (left) and $C_{Pet20}$-motifs (right) of Pet20 (A) and Sue1 (B) colored based on their physico-chemical properties (color intensity corresponds to level of conservation) and the IPR014804/PF08692 consensus. Residues predicted for hydrogen bond formation with the Pim1 NTD are indicated with black triangles. Blue triangles above $C_{Pet20}$-motif of Pet20 indicate that these residues are predicted to form hydrogen bonds to another region of the Pim1 NTD in predictions where the $N_{Pet20}$-motif binds the Pim1 NTD interaction site (see also as blue line in C). (**C**) Pairs of residues predicted for hydrogen bond formations between Pet20 proteins and the Pim1/LONP1 NTD in highest ranked models where either the $N_{Pet20}$- or $C_{Pet20}$-motif interacts with the Lon protease (see Supplementary Figure S4B). Amino acids are colored based on their physico-chemical properties. Aliphatic/ hydrophobic residues (ILVAM) are colored in salmon, aromatic residues (FWY) in orange, positive charged residues (KRH) in blue, negative charged residues (DE) in red, hydrophilic residues (STNQ) in green, structurally distinctive residues (PG) in magenta and cysteines (C) in yellow. Black lines next to residues indicate that these residues are not positioned within the conserved Pet20 motif regions. (**D**) Consensus logo of 34 distantly related eukaryotic Lon protease homologue NTDs reviewed from IPR027503. Numbering of the aligned residues refers to the amino acid sequence of *S. cerevisiae* W303 Pim1. Positions of residues involved in hydrogen bond formation to Pet20 domain proteins are marked with geometric forms as indicated. Residues are colored based on their physico-chemical properties as in (C). Hydrophobic stretches implicated in interaction with the Mrx6 Pet20 motifs inferred from S7F are highlighted on the bottom by salmon triangles where different contours specify the type of interaction. No contour: $N_{Pet20}$- motif, blue contour: hydrophobic $N_{Pet20}$-motif link to next adjacent Pim1 NTD, black countour: $C_{Pet20}$-motif. (**E**) MUSCLE (Edgar, 2004) protein alignment of 34 distantly related eukaryotic Lon protease homologue NTDs reviewed from IPR027503 with extracted fungi and animalia species as indicated. Residues are colored based on their physico-chemical properties as in (C) . A conservation threshold of 30 was applied on the color opacity in Jalview (Waterhouse et al., 2009). Numbering on top of the aligned residues refer to the amino acid sequence of *S. cerevisiae* W303 Pim1.

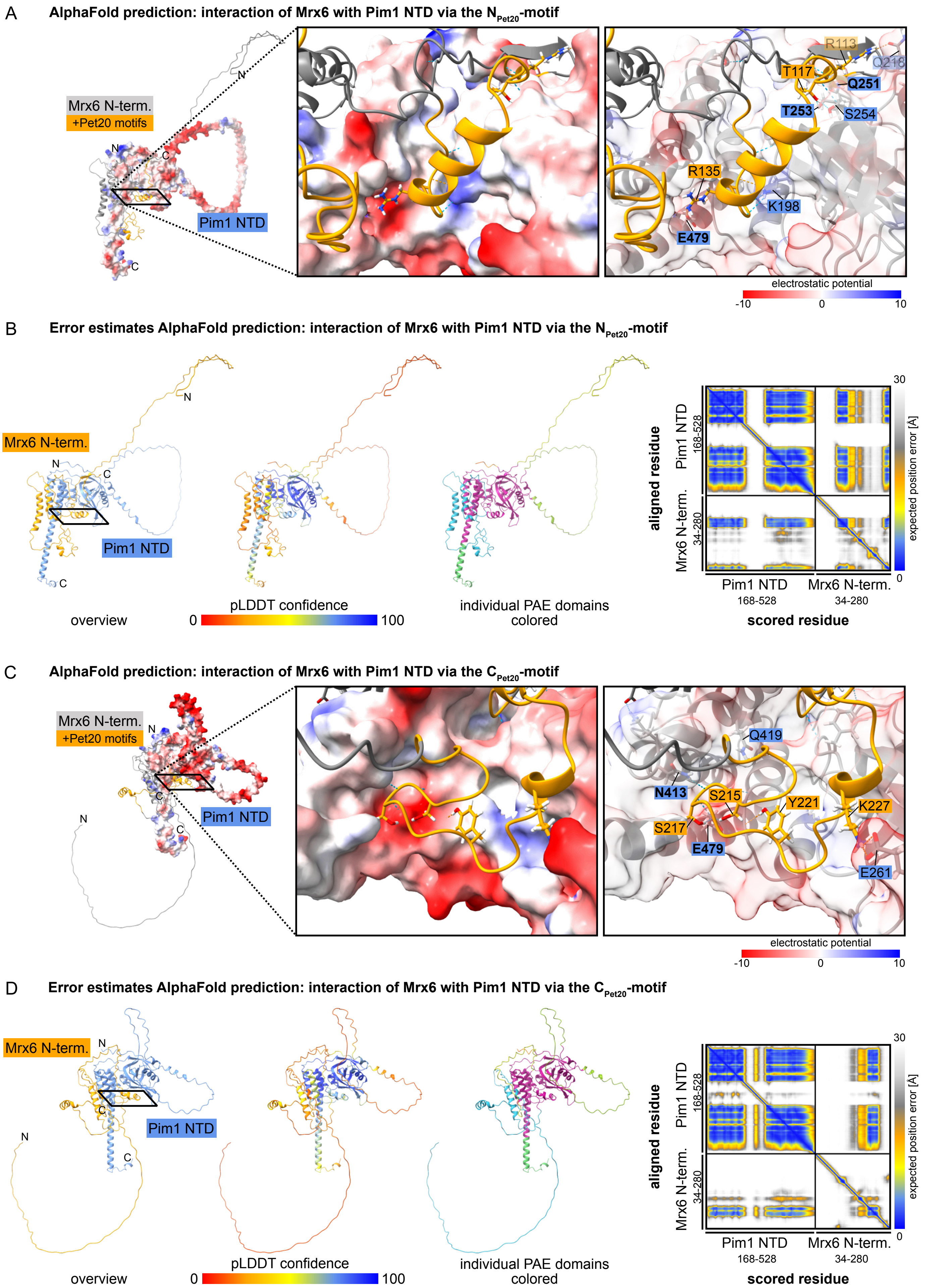


**Figure S6. Mrx6 is predicted to interact with the Pim1 NTD either via the $N_{Pet20}$-motif or the $C_{Pet20}$-motif, while the predicted human LONP1 interaction with Mrx6 is restricted to the $C_{Pet20}$-motif.**
Related to Figure 3. see full legend on next page.

E **AlphaFold prediction: interaction of Mrx6 with LONP1 NTD via the $C_{Pet20}$-motif**

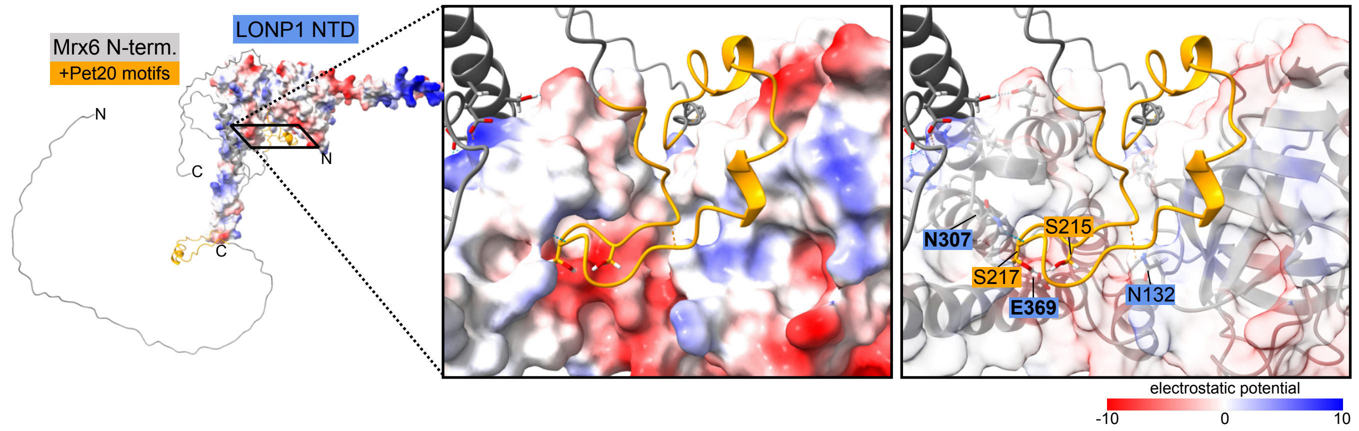


F **Error estimates AlphaFold prediction: interaction of Mrx6 with LONP1 NTD via the $C_{Pet20}$-motif**

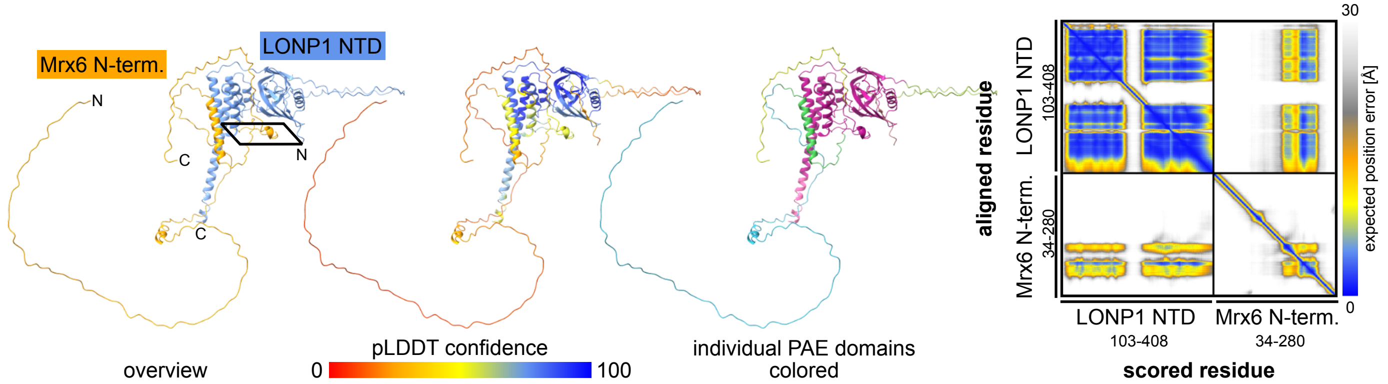


**Figure S6. Mrx6 is predicted to interact with the Pim1 NTD either via the $N_{Pet20}$-motif or the $C_{Pet20}$-motif, while the predicted human LONP1 interaction with Mrx6 is restricted to the $C_{Pet20}$-motif.**
Related to Figure 3.

(**A-F**) AlphaFold multimer predictions of Mrx6 N-terminus (residues 34-280) together with the Pim1 NTD (residues 168-528) either in a conformation, where the $N_{Pet20}$-motif interacts with the Pim1 NTD (best ranked model with ipTM + pTM confidence of ~0.6582) (**A+B**), or the $C_{Pet20}$-motif interacts with the Pim1 NTD (best ranked model with ipTM + pTM confidence of ~0.6648) (**C+D**). The interaction of the Mrx6 N-terminus (residues 34-280) with the LONP1 NTD (residues 103-408) is predicted only in a state where the $C_{Pet20}$-motif interacts with LONP1 (best ranked model with ipTM + pTM confidence of ~0.6823) (**E+F**). (**A, C, E**) Detailed view of the respective predicted interaction sites and involved residues in hydrogen bond formation with a full size overview structure on the left where the position of the zoom is indicated by a black rectangle. The Mrx6 N-terminus is shown as gray ribbon with Pet20 motifs highlighted in orange. The Pim1/LONP1 NTD is shown as surface structure colored according to the coulombic electrostatic potential in the range of -10 to 10 with full opacity on the left and the middle views or with transparency on the right with a gray colored ribbon. Labels for critically involved Pim1/LONP1 NTD residues are colored blue. (**B, D, F**) Error estimates of the respective predictions are shown next to an overview depiction with the Mrx6 N-terminus colored in orange and the Pim1/LONP1 NTD in blue. Error estimates are colored in the ribbon structures according to the residue specific pLDDT confidence or coherent domains are colored separately based on residue clustering according to the predicted aligned (PAE) value, additionally depicted as a heat map plot of the expected position error for every pair of residues.

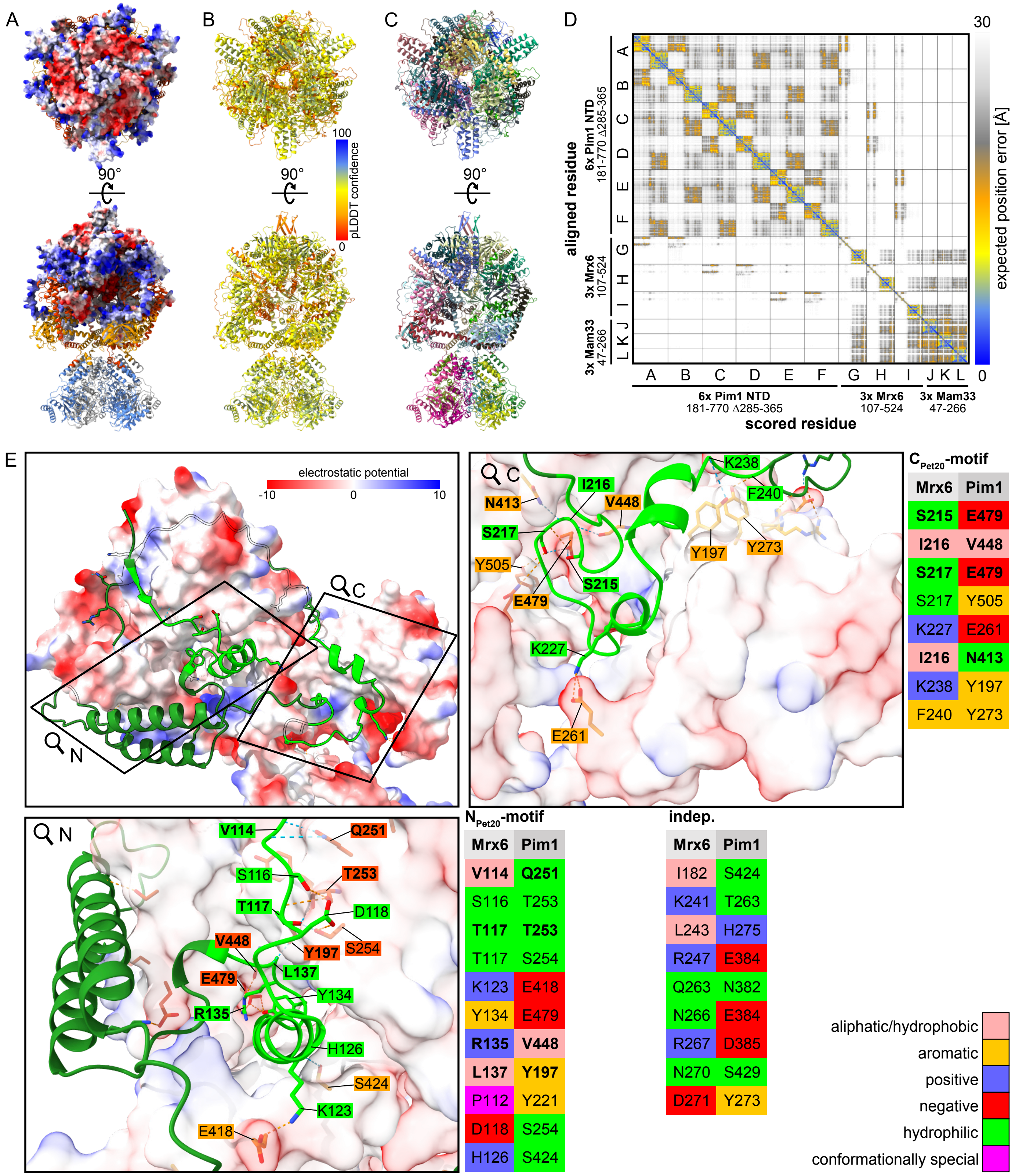


**Figure S7. A bipartite-motif in the Pet20 domain mediates interaction between Mrx6 and Pim1.** Related to Figure 3.
AlphaFold 3 multimer prediction of 6x Pim1 NTD + ATPase (181-770 with deleted disordered region 285-365), 3x Mrx6 (107-524) and 3x Mam33 (47-266) best ranked model with ipTM + pTM confidence of ~0.46. (**A**) Ribbon structure shows the Pim1 protomers colored separately for the N-terminal domains (181-528) (orange/orange-red) or the ATPase domains (529-770) (gray/blue) in an alternating manner (see labels Figure 3C). 3x Mrx6 and 3x Mam33 are depicted as surface structures colored according to the coulombic electrostatic potential in the range of -10 to 10. (**B-D**) Error estimates shown as ribbon structure colored according to the residue specific pLDDT confidence (B), separate coloring of coherent domains based on residue clustering according to the predicted aligned (PAE) value (C), or depicted as a heat map plot of the expected position error for every pair of residues (D). (**E**) Detailed view of the interaction between the Mrx6 N-terminus and Pim1 NTD chains B (left, acute-angled) and A (right, obtuse-angled) via the $N_{Pet20}$- and $C_{Pet20}$-motif, respecitvely. Overview shows same view as Figure 3F. The Mrx6 N-terminus ribbon is colored in dark green and Pet20 motifs in lime. The two Pim1 NTDs are shown as surface structure colored according to the coulombic electrostatic potential in the range of -10 to 10. The position of the zoom for the detailed view of involved hydrogen bonds for the interaction between either the $N_{Pet20}$- or $C_{Pet20}$-motif with either the acute- or obtuse-angled Pim1 NTD is indicated by a black rectangle. Labels for residues involved in hydrogen bond formation are colored in lime (bipartite Pet20 motif of Mrx6), orange-red (acute-angled Pim1 NTD chain B) or orange (obtuse-angled Pim1 NTD chain A). Pairs of hydrogen bonds inferred from the entire structure of hexameric Pim1 and trimeric Mrx6 consistently predicted for all three Mrx6 subunits each interacting with one of the three different pairs adjacent acute- and obtuse-angled Pim1 NTDs are listed. Pairs are separated based on the involvement of the $N_{Pet20}$- or $C_{Pet20}$-motif or additional Pet20 domain independent residues of Mrx6. The background of the single letter amino acid code is colored based on their physico-chemical properties. ...

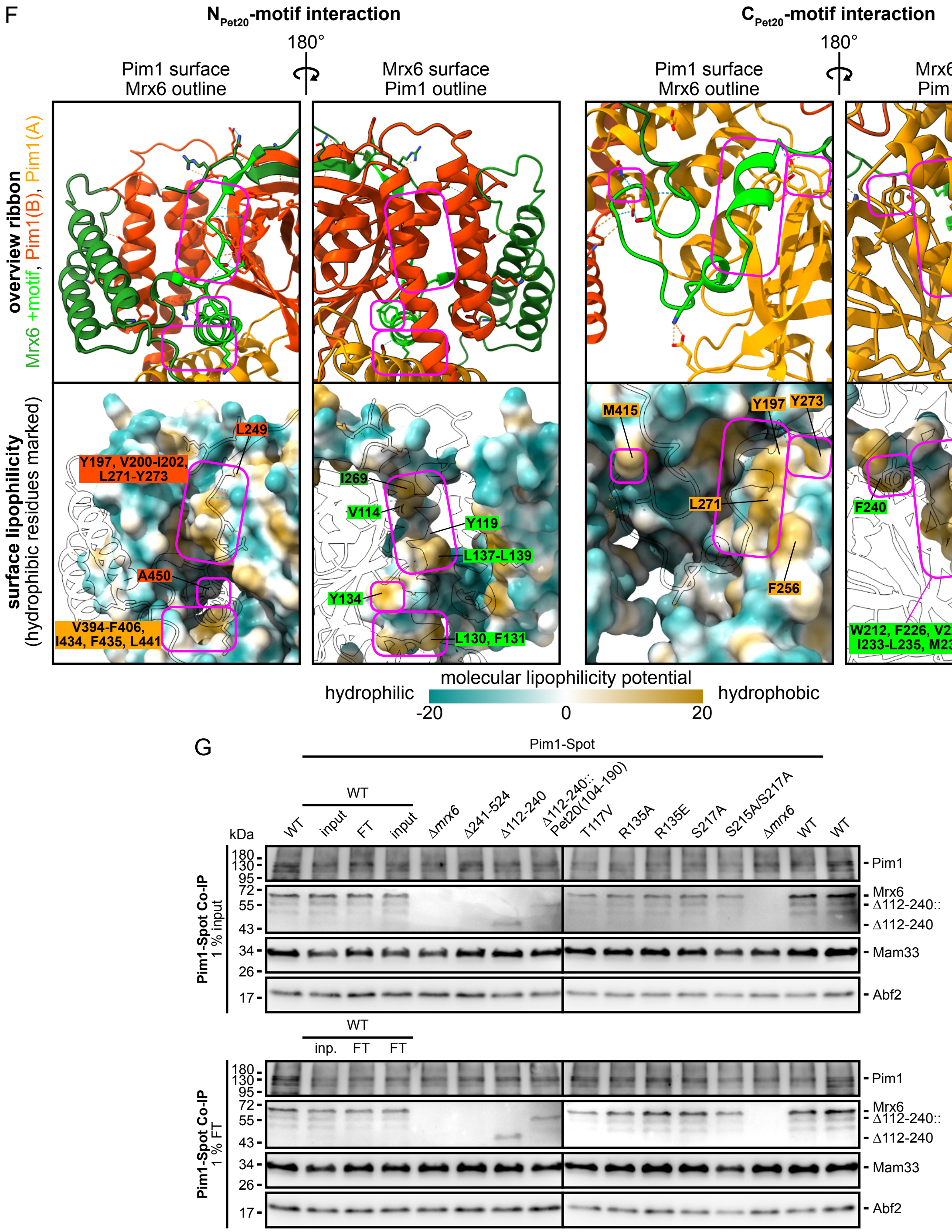


**... Figure S7 continued** (**F**) Detailed view of hydrophobic surface interactions between the Mrx6 $N_{Pet20}$- (left) and $C_{Pet20}$-motifs (right) and Pim1 NTD chains B (acute-angled), and A (obtuse-angled). Views are rotated around the y-axis by 180° to reveal the direct contact of hydrophobic surfaces between Pim1 (left) and Mrx6 (right). Ribbon overviews in upper panel show the Mrx6 N-terminus colored in dark green with lime Pet20 motifs and the two Pim1 NTDs in orange-red (acute-angled, B) or orange (obtuse-angled, A). In lower panels the ribbon of one of the interacting protein is only shown by a black outline of the ribbon while the other one is depicted as surface structure colored according to the molecular lipophilicity potential in the range of -20 to 20. Labels for residues are colored like the respective ribbons when they are involved in the hydrophobic contacts, which are highlighted with magenta rounded rectangles. (**G**) Immunoblots of Pim1, Mrx6, Mam33 and Abf2 (control) derived from 1% input and flow-through (FT) control fractions of native total cell lysate immunoprecipitations of Pim1-Spot from indicated strains cultivated in YPG at 30°C shown in Figure 3I. Δ112-240 and Δ241-524:: mark running heights of the Mrx6 versions with deleted Pet20 domain or its replacement with corresponding residues 104-190 from Pet20. C.B.B.: colloidal Coomassie stained high molecular weight proteins that retained in the gel after transfer.

- E479 cavity of Pim1: E479 is surrounded by N482, A450, Q486, N413, I409, A483, F416, V448 and Q419
- Further hydrogen bond interaction sites: The hydrogen of the amide in the side chain of Pim1 Q251 is a hydrogen bond donor for the peptide bond carbonyl oxygen of Mrx6 V114, Pet20 V106 or Sue1 V61. In Mrx6, the secondary amine of the V114 peptide bond donates a hydrogen bond to the oxygen of the amide group in the side chain of Pim1 Q251. The same Pet20 motif residues form additional hydrogen bonds, with their backbone carbonyl oxygen accepting a hydrogen bond from the hydrogen of the amide group in the side chain of Pim1 N413.
- Additional stabilizing hydrogen bonds of the bipartite motif in the oligomeric prediction: The juxtaposition of both Pim1 NTDs is stabilized by the positive charged residues K123 and H126 of the $N_{Pet20}$-motif that bridge to residues E418 and S424 of the proximate obtuse-angled NTD, which binds the $C_{Pet20}$-motif. Independent of the bipartite motif, an additional interaction with the outward facing part of the acute-angled NTD is achieved by threading around Mrx6 residues 241-271, which are placed between the $C_{Pet20}$-motif and the Mrx6- specific C-extension.

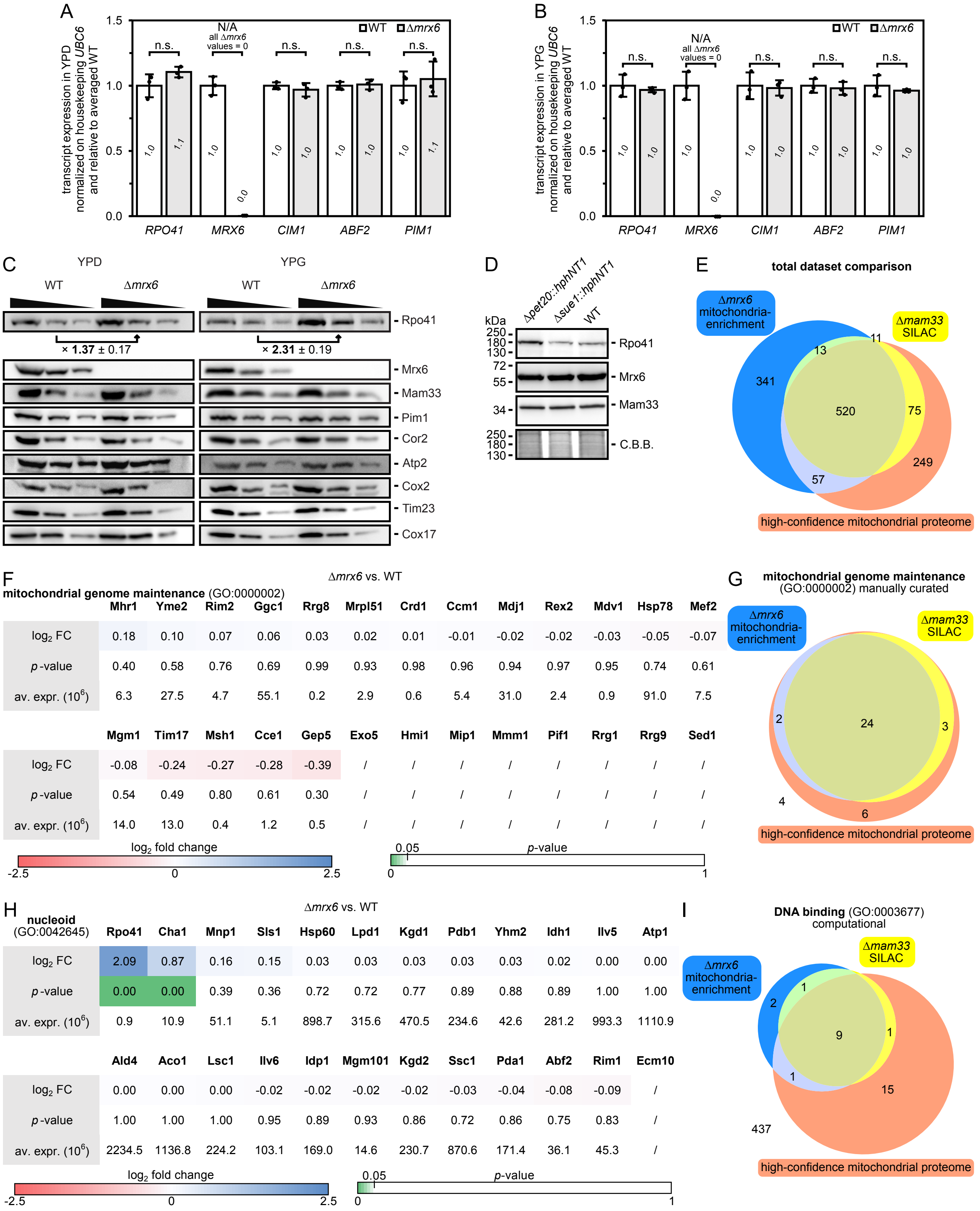


**Figure S8. Absence of Mrx6 leads to a post-transcriptional increase of Rpo41 and Cim1.**

Related to Figure 4.

(**A+B**) RT-qPCR transcript comparison of indicated genes between WT and Δ*mrx6* with RNA from cells cultivated in YPD (A) or YPG (B) medium at 30°C. Normalization was performed to housekeeping gene *UBC6* and relative to the averaged WT. n = 3 biological replicates, data represent mean ± SD.(**C**) Immunoblots of indicated mitochondrial proteins, including mtDNA encoded Cox2, of total cell lysates in serial 1:2 dilutions from WT and Δ*mrx6* cells cultivated in YPD or YPG at 30°C. Rpo41 intensities were quantified from 3 biological replicates normalized to the respective WT, data represent mean ± SD. (**D**) Immunoblots of Rpo41, Mrx6 and Mam33 of lysates from indicated cells grown in YPG at 30°C. Loading was additionally controlled by high molecular weight C.B.B. stained protein bands that retained in the gel after transfer. n = 1. (**E**) Total data set comparison between LFQ LC-MS/MS derived data from mitochondrial enriched isolates of Δ*mrx6* vs. WT cells, a SILAC dataset filtered for mitochondrial proteins of Δ*mam33* vs. WT cells (Hillman & Henry, 2019) and a high-confidence mitochondrial proteome (Morgenstern et al., 2017). (**F**) Δ*mrx6* vs. WT dataset $\log_2$ fold changes, p-values and average expression of detected mitochondrial genome maintenance proteins described by GO:0000002. (**G**) Comparison of GO:0000002 coverage between indicated datasets. (**H**) Δ*mrx6* vs. WT dataset $\log_2$ fold changes, p-values and average expression of detected mitochondrial nucleoid proteins described by GO:0042645. (**I**) Comparison of GO:0003677 (annotated for DNA binding) coverage between indicated datasets.

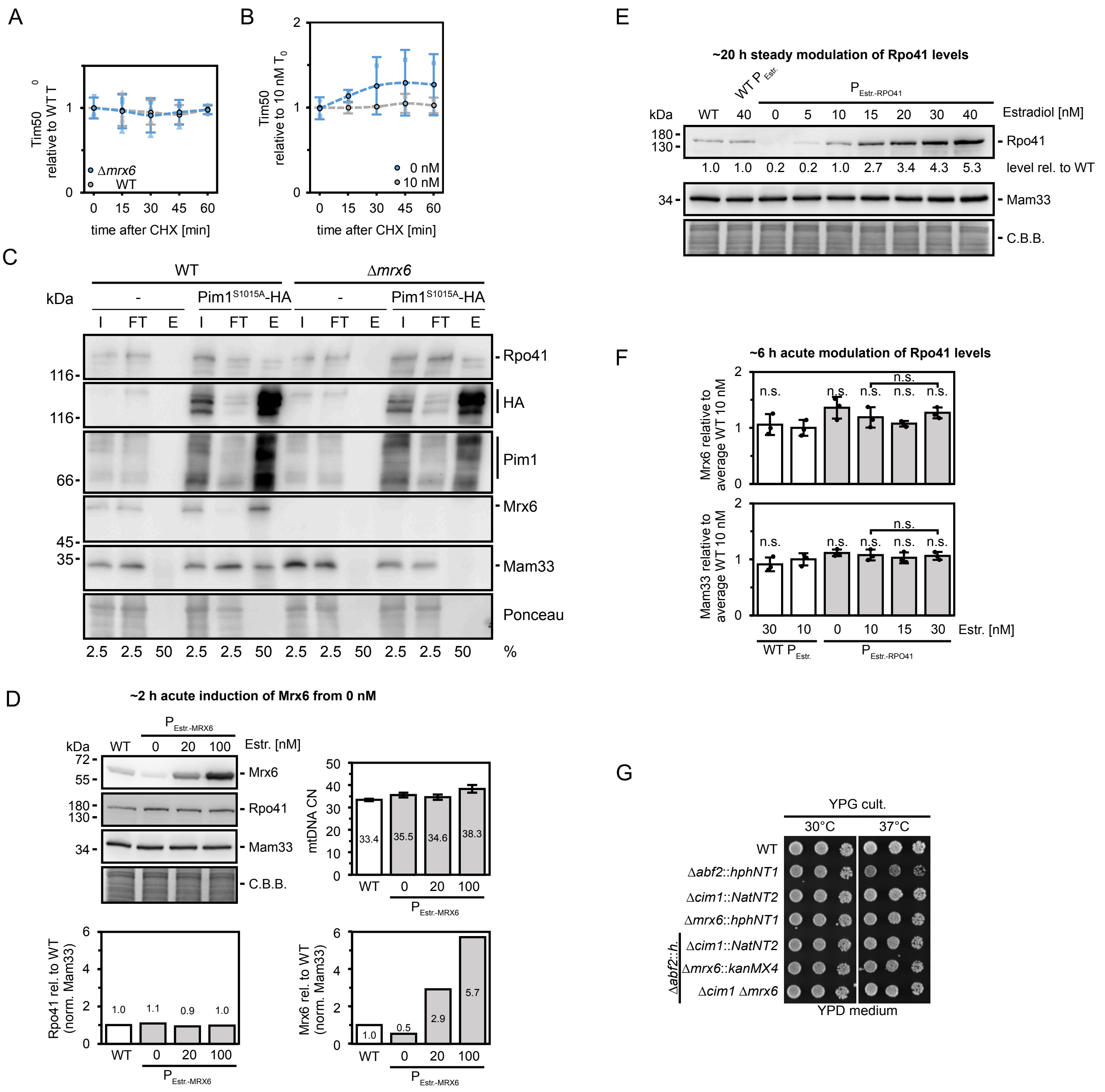


**Figure S9. Alterations of the Pim1-Mrx6 complex stabilize multiple factors involved in the regulation of the mitochondrial genome.**
Related to Figures 5 and 6.

(**A**) Quantification of Tim50 (loading control) immunoblots of lysates from WT and Δ*mrx6* cells cultivated in log phase in YPG at 30°C and chased for one hour after addition of cycloheximide (CHX). n = 3, data represent mean (dots with black outline) ± SD. (**B**) Quantification of Tim50 (loading control) immunoblots of lysates from $P_{Estr.}$-*PIM1* cells cultivated in log phase in YPG at 30°C and chased for one hour after addition of CHX. WT-like Pim1 levels were sustained with 10 nM estradiol, whereas prior withdrawal of estradiol for ~18 h resulted in downregulation. n = 2, data represent mean (dots with black outline) ± SD. (**C**) Immunoblots with indicated antibodies from HA-tag immunprecipiations of lysates from WT or Δ*mrx6* cells containing empty vector or a construct expressing $Pim1^{S1016A}$-HA. Percentage of loaded sample is shown. Ponceau staining was used as a loading control. (**D**) Immunoblots of Mrx6, Rpo41 and Mam33 (loading control) of lysates from $P_{Estr.}$-*MRX6* cells grown for ~2 h in YPG at 30°C in the presence of 0, 20 or 100 nM estradiol, which in $P_{Estr.}$-*MRX6* strains results in reduced, WT-like or overexpressed levels of Mrx6. WT cells without expression of any components of the estradiol system served as control. C.B.B.: colloidal Coomassie stained high molecular weight proteins that retained in the gel after transfer. Mrx6 and Rpo41 signals were quantified relative to WT and further normalized on Mam33 signals. mtDNA CN of the same cell samples was determined by qPCR on mtDNA encoded *COX1* and nuclear reference *TAF10*. n = 1, data represent technically derived mean ± SD including error propagation of pipetting replicates of DNA samples. (**E**) Immunoblots of Rpo41 and Mam33 (loading control) of lysates from $P_{Estr.}$-*RPO41* cells grown for ~20 h in YPG at 30°C in the absence, or presence of increasing estradiol concentrations, starting from cell cultures that have been cultivated in the presence of 10 nM estradiol. Cells with WT *RPO41* promoter lacking LexA binding sites served as control. In these cells either the LexA-(estradiol receptor ligand binding domain)-B42 activator fusion construct was present and 40 nM estradiol was added to the medium (WT $P_{Estr.}$) or absent (WT). Rpo41 signals were quantified relative to WT. C.B.B.: colloidal Coomassie stained high molecular weight proteins that retained in the gel after transfer. n = 1. (**F**) Quantification of immunoblots shown in Figure 6A. (**G**) Drop dilution growth analysis at 30°C or 37°C of indicated strains pre-grown in YPG at 30°C. Images were taken after 48 h growth on YPD. Only dilution steps 2-4 are shown.
